# Steric bulk at one position of a tight junction modulator switches the kinetic step governing angulin paralog selectivity

**DOI:** 10.64898/2026.11.26.769419

**Authors:** Taiki Kuzu, Yuki Niwa, Shizuki Kimura, Masuo Kondoh

## Abstract

Three angulin paralogs organize tricellular tight junctions, which control paracellular macromolecular passage. No angulin–ligand complex structure exists, and how ligands discriminate these paralogs is unknown. Using AlphaFold3 to nominate the interface, we defined a 120-residue scaffold, angubindin-2, retaining its parent angubindin-1’s dual angulin-1/angulin-3 binding and reversible barrier modulation. A saturation scan ranked Ala14 first for predicted discrimination. Ala14 substitutions shifted preference in both directions: threonine and valine abolished detectable angulin-3 binding, increasing selectivity from 31-fold to above 370-fold, whereas glycine increased angulin-3 affinity sevenfold. Surface plasmon resonance showed where side-chain bulk acts: at angulin-3, the penalty was complete by the transition state, slowing association 75-fold; at angulin-1, it appeared only in the complex, shortening its lifetime 13-fold. A14T and A14V reduced transepithelial resistance through angulin-1 but not angulin-2 or angulin-3. One position therefore sets the receptor preference and the kinetic step that carries it, making selectivity and duration separable.

## Introduction

Tight junctions (TJs) seal the paracellular space and define the selective permeability of epithelial and endothelial barriers^1,2^. Structurally and functionally, TJs have two compartments: bicellular tight junctions (bTJs), at contacts between two cells, and tricellular tight junctions (tTJs), at the convergence of three cells. Tricellulin is a barrier-forming component of tricellular contacts^3^, and the angulin family proteins angulin-1/lipolysis-stimulated lipoprotein receptor (LSR), angulin-2/immunoglobulin-like domain-containing receptor 1 (ILDR1), and angulin-3/ILDR2 organize tTJs, marking cell corners and recruiting tricellulin^4,5^. Because tTJs regulate the passage of macromolecules largely excluded by bTJs^6^, tTJs are relevant both to barrier integrity in tissues including the blood–brain barrier^7^ and inner ear^8^, and to paracellular delivery across epithelial and endothelial barriers.

Binary iota-family toxins use angulin-1 as their host receptor^9,10^, and their receptor-binding region is the principal molecular tool used to investigate this system. Angubindin-1 (Ang1), derived from the C-terminal receptor-binding region of the *Clostridium perfringens* iota-toxin Ib component, binds angulin-1 and angulin-3, reversibly modulates paracellular barrier function, and enhances macromolecular permeation across epithelial^11^ and endothelial^12^ barriers. Alanine-scanning mutagenesis of Ang1 (Ib421–664) identified six residues — L562, L598, E638, V640, Y643, and K644 — required for angulin binding and tTJ modulation^13^, while comparisons of toxin-derived receptor-binding domains have begun to define their permeation properties^14^. Structural understanding has progressed unevenly between these two compartments. At bTJs, structural information has made the equivalent problem tractable: the C-terminal fragment of *Clostridium perfringens* enterotoxin (C-CPE) both removes specific claudins from TJ strands^15^ and has been resolved in complex with its claudin receptors, defining the interface residues that set subtype preference^16,17^. No experimentally determined structure of an angulin family protein bound to a toxin-derived ligand exists, leaving the local features that distinguish the closely related paralogs angulin-1 and angulin-3 unresolved.

Loss-of-function mapping, however, cannot resolve how selectivity between paralogous receptors is encoded because residues essential for binding one receptor are frequently essential for binding all receptors. Moreover, Ang1’s engagement of both angulin-1 and angulin-3 prevents separation of angulin-1-specific effects from those of broader angulin engagement. The converse experiment is required instead: to locate, among the residues shared by both receptors, the position at which the two interfaces diverge, and to test substitutions there for a gain of selectivity. Without a structure of either complex, that position cannot be read from coordinates.

Predicted complexes can supply that position as a hypothesis. AlphaFold3 (AF3) predicts protein–protein complexes without requiring an experimental assembly template^18^ and provides testable structural hypotheses in the absence of experimental structures. Its interface confidence metrics, however, describe predicted models rather than binding affinity, and predicted-structure confidence responds only weakly to single amino-acid substitutions^19^. Whether systematic *in silico* mutagenesis at a predicted interface can identify in advance a position that shifts preference between closely related receptor paralogs, and whether substitutions at that position confer receptor-dependent biological function, remain open questions. Prediction might therefore be used to nominate where to look, leaving protein-level binding and function to establish what to put there — a division of labor distinct from *de novo* binder design, which begins from a target structure^20,21^.

We tested this division of labor in the angulin system. Using AF3, we modeled Ang1 in complex with each angulin family member and identified a compact, interface-containing ligand region, expressed as a self-contained construct designated angubindin-2 (Ang2). By retaining Ang1’s receptor-binding and barrier-modulating activity, Ang2 provides a minimal scaffold for systematically probing the angulin-binding surface. The *in silico* scan nominated one specificity-determining position, and saturation mutagenesis there, comparing all 20 amino acids under identical computational and experimental conditions, resolved outcomes that prediction could not distinguish: flow cytometry (FCM) identified four that retained angulin-1 recognition while losing detectable angulin-3 binding, of which two retained barrier-modulating activity through angulin-1 alone. Binding kinetics explained the receptors’ different responses to the same change: each receptor incurred the penalty at a different step in the binding reaction.

These results show that discrimination between closely related angulin paralogs can be shifted by a single residue within a minimal binding scaffold. The principal finding is kinetic: the number of side-chain atoms beyond Cβ at that position determines which step of the binding reaction carries the cost of substitution, and that step differs between the receptors — association at angulin-3, complex lifetime at angulin-1. Selectivity at this interface is therefore set by where in the binding reaction a substitution is felt rather than by the magnitude of its effect, and the same residue supplies two separable handles: the height of the angulin-3 association barrier and the depth of the angulin-1 well. This mechanism and the selective mutants it yields together provide a framework for engineering paralog-selective modulators of protein–protein interactions lacking experimental structures.

## Results

### AlphaFold3 (AF3) identifies a minimal angulin-binding scaffold within angubindin-1 (Ang1)

Because no Ang1–angulin complex structure is available, we generated interface hypotheses with AF3, using the models to nominate residues for experimental testing rather than treating them as experimentally determined structures. AF3 models of Ang1 in complex with the human angulin-1 extracellular domain (ECD) predicted two distinct regions joined by an extended, low-confidence segment (Figure 1A). In top-ranked models, all predicted intermolecular contacts with the angulin-1 ECD lay within the C-terminal region spanning Ib545–664; none involved the N-terminal region (Ib421–544) (Figure 1A). We therefore designed a domain-optimized scaffold comprising Ib545–664, designated angubindin-2 (Ang2). We set the boundaries to retain the entire predicted receptor interface and exclude regions predicted to lie outside it, using more than per-residue confidence alone (Figure 1B). Re-prediction of Ang2 against all three human angulin ECDs reproduced the parent molecule’s paralog pattern: high interface confidence for angulin-1 (ipTM 0.86, pTM 0.80) and angulin-3 (ipTM 0.86, pTM 0.79), and low interface confidence for angulin-2 (ipTM 0.19, pTM 0.57) (Figure 1C–1E).

**Figure 1.**
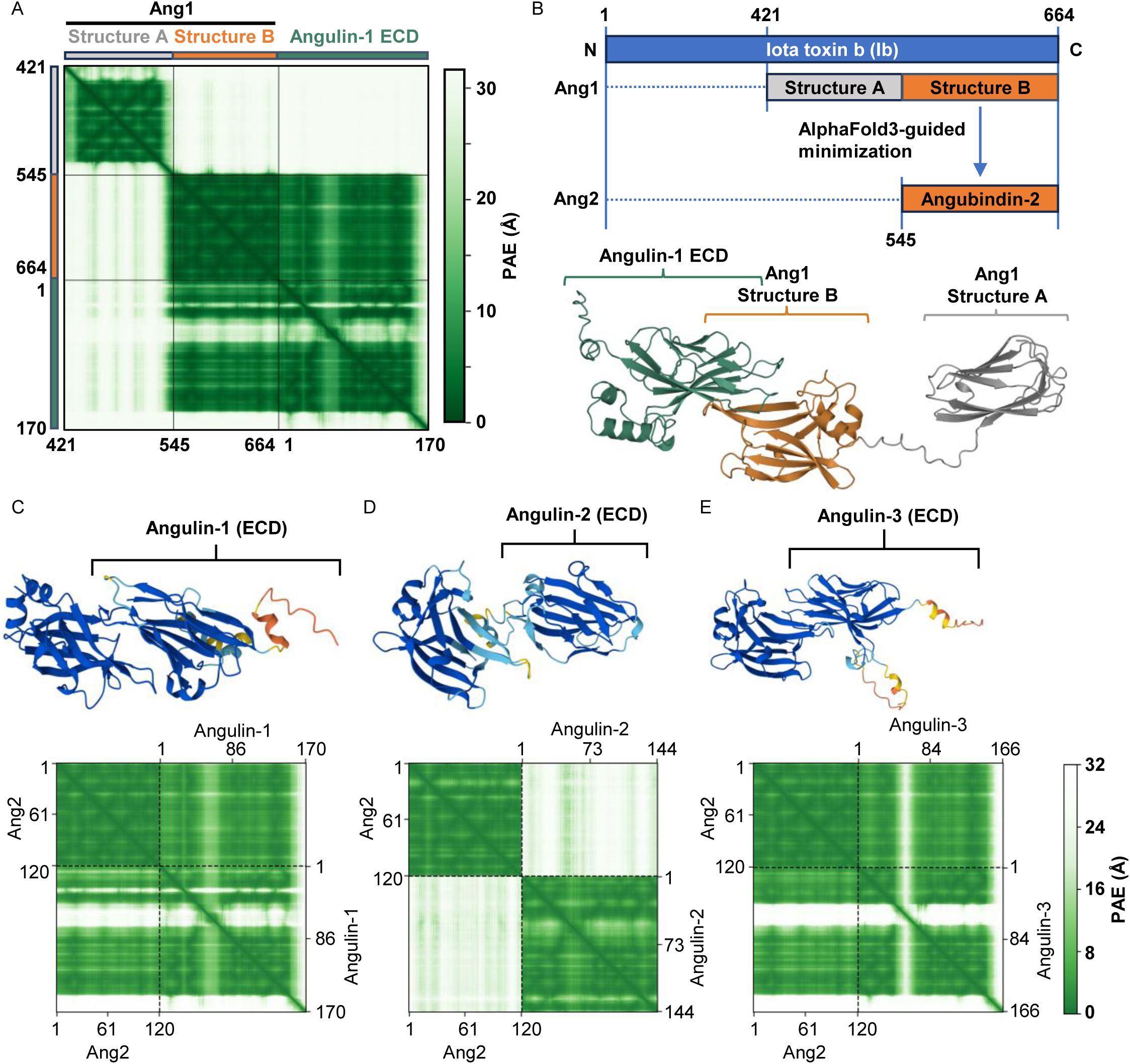
AF3 models of Ang1–angulin complexes define a minimal angulin-binding scaffold, Ang2. Models were generated with AF3 using *Clostridium perfringens* iota-toxin receptor-binding component Ib as ligand and each human angulin ECD as receptor; Ang1 numbering follows mature Ib (Ib421–664). (A) Predicted aligned error (PAE) matrix of the top-ranked Ang1 (Ib421–664)–angulin-1 ECD (residues 1–170) model. Residues are numbered 1–170 according to the construct, corresponding to residues 42–211 of angulin-1 (NP_991403.2). Ang1 contains two predicted regions, Structure A (Ib421–544) and Structure B (Ib545–664), joined by an extended low-confidence segment. Off-diagonal confidence with angulin-1 is confined to Structure B; Structure A has no predicted receptor contact. (B) Top, domain organization of Ib (residues 1–664), Ang1 (Ib421–664; Structure A, gray; Structure B, orange), and AF3-guided minimized scaffold Ang2 (Ib545–664, 120 aa). Bottom, top-ranked Ang1–angulin-1 ECD ribbon model, colored as above (angulin-1, green), showing Structure B engaging the receptor and Structure A away from the interface. (C–E) AF3 models of Ang2 with angulin-1 (C), angulin-2 (D), and angulin-3 (E) ECDs. Top, receptor ECD ribbons; bottom, corresponding complex PAE matrices, with the PAE scale (0–32 Å) at right. Ang2 is numbered 1–120 according to the construct, corresponding to Ib545–664. Residues of receptors are numbered according to each construct: angulin-1, 1–170 (residues 42–211 of NP_991403.2); angulin-2, 1–144 (residues 24–167 of NP_001186728.1); angulin-3, 1–166 (residues 21–186 of NP_955383.1). Top-ranked model interface/whole-complex confidence values were ipTM 0.86/pTM 0.80 for angulin-1, ipTM 0.19/pTM 0.57 for angulin-2, and ipTM 0.86/pTM 0.79 for angulin-3, reproducing Ang1’s paralog preference. Each single-seed job returned five diffusion samples, ranked within the job by ranking score.

### Ang2 retains angulin binding and tTJ-modulating activity

We next tested whether Ang2 retains full scaffold function. Binding was measured by FCM using parental HT1080 cells and cells expressing angulin-1, angulin-2, or angulin-3. Ang2 and Ang1 showed the same binding profile: both proteins bound HT1080/angulin-1 and HT1080/angulin-3 cells, whereas binding to HT1080/angulin-2 cells did not exceed parental background for either protein under the tested conditions (Figure 2A). We also compared binding in Caco-2 cells^22^, the line used for the functional assays below, to assess binding and barrier modulation in the same cellular context. The extent of Caco-2 cell binding was comparable for Ang1 and Ang2 (Figure 2B), indicating that the amount engaging the cell surface is unaffected by their difference in size. Barrier modulation was assessed in Caco-2 monolayers, in which Ang1 was originally defined as a tTJ modulator. To account for their different sizes, proteins were compared on an equimolar rather than mass basis, at 0.1, 0.3 and 1 µM for 24 h. At 0.1 µM, neither protein altered barrier function: transepithelial electrical resistance (TER) at 24 h was 332.6 Ω·cm^2^ for the vehicle control, 328.5 Ω·cm^2^ for Ang1 and 359.2 Ω·cm^2^ for Ang2, with neither protein differing significantly from vehicle. Both proteins produced comparable TER reductions at 0.3 and 1 µM (0.3 µM: 62% of vehicle for Ang1 vs. 67% for Ang2; 1 µM: 49% for Ang1 vs. 57% for Ang2); neither concentration produced a significant difference between the proteins (Figure 2C). We tested reversibility by washing out both proteins after 24 h of treatment and monitoring TER for a further 24 h. TER returned to vehicle levels by 48 h at every concentration tested, for both Ang1 and Ang2 (Figure 2D), indicating that the decrease in TER reflects transient, reversible barrier opening rather than cytotoxicity or irreversible damage to the monolayer.

**Figure 2.**
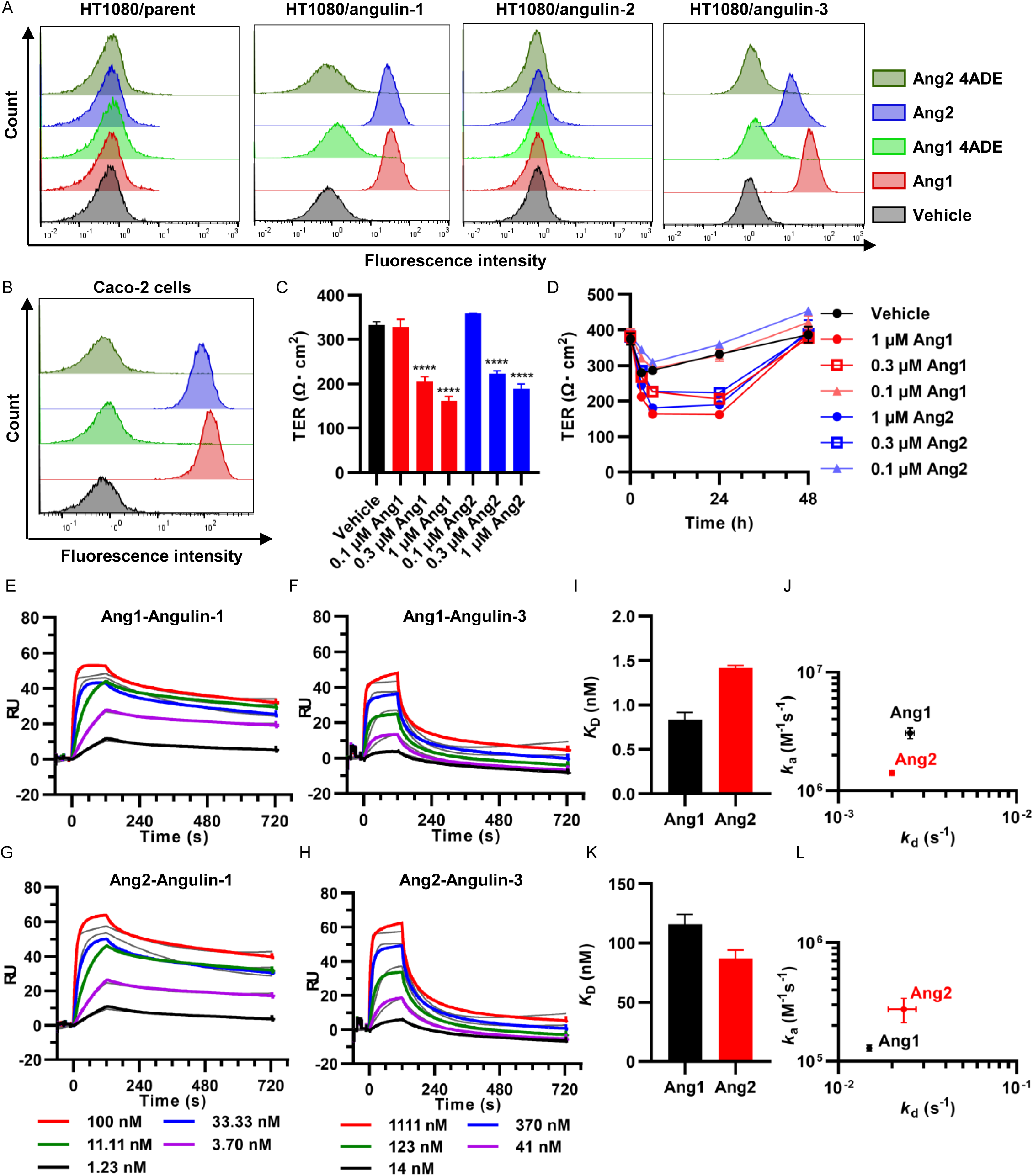
Ang2 retains Ang1’s angulin-binding profile, barrier-modulating activity, and binding kinetics. (A) FCM analysis of protein binding to parental HT1080 cells and cells stably expressing FLAG-tagged angulin-1, angulin-2, or angulin-3. Cells received 1 µM Ang1, interface-null Ang1 4ADE, Ang2, or Ang2 4ADE, or vehicle. Bound protein was detected using an anti-His-tag antibody, biotin-SP-conjugated anti-mouse IgG, and Alexa Fluor 647-conjugated streptavidin. Representative histograms from three independent experiments are shown. Ang1 and Ang2 bound angulin-1- and angulin-3-expressing cells but showed no signal above parental background on angulin-2-expressing cells; 4ADE controls remained at background throughout. (B) FCM analysis showing comparable Ang1/Ang2 binding to Caco-2 cells used for functional assays. (C) Caco-2 monolayer TER 24 h after basal addition of Ang1 or Ang2 at 0.1, 0.3, or 1 µM. Proteins were compared equimolarly. Neither altered TER at 0.1 µM; both reduced it comparably at 0.3 and 1 µM. (D) Reversibility: proteins were washed out after 24 h, and TER was followed for another 24 h. Both proteins yielded recovery to vehicle levels by 48 h at every concentration. (E–H) SPR sensorgrams with His-tagged Ang1 or Ang2 captured on NTA chips and angulin ECD–rabbit Fc analytes: Ang1–angulin-1 (E), Ang1–angulin-3 (F), Ang2–angulin-1 (G), and Ang2–angulin-3 (H). Colored traces are double-referenced responses at concentrations indicated below panels (1.23–100 nM for angulin-1; 14–1111 nM for angulin-3); gray traces are global 1:1 Langmuir fits. (I, J) Angulin-1 equilibrium dissociation constant *K*_D_ (I) and association rate constant *k*_a_ vs. dissociation rate constant *k*_d_ (J). (K, L) Angulin-3 *K*_D_ (K) and *k*_a_ vs. *k*_d_ (L). Both proteins bound angulin-3 60- to 140-fold more weakly than angulin-1 through different kinetic mechanisms. In (I–L), bars and symbols are arithmetic means ± SEM from three independent SPR experiments. In (C) and (D), bars are means ± SEM of replicate wells within a single experiment, each well a separate insert carrying its own monolayer (*n* = 3 wells per group), and the experiment shown is representative of at least three independent repeats. Significance was assessed by one-way ANOVA with Dunnett’s multiple-comparison test against vehicle: \*\*\*\**P* < 0.0001.

We quantified direct binding by surface plasmon resonance (SPR), capturing Ang1 or Ang2 on the sensor surface and using angulin ECD–rabbit Fc fusion proteins as analytes (Figure 2E–2H). Ang2 bound both receptors with affinities within 2-fold of those of Ang1 (angulin-1, 1.42 vs. 0.84 nM; angulin-3, 87.3 vs. 116.0 nM), although the underlying rate constants differed: relative to Ang1, Ang2 associated approximately 2-fold more slowly with angulin-1 and approximately 2-fold more rapidly with angulin-3, with smaller differences in dissociation (Supplementary Tables 1 and 2, Figure 2I–2L). Ang2 thus preserves dual angulin binding and tTJ-modulating activity in a minimal scaffold. The kinetics indicate separable determinants of angulin-1 and angulin-3 interactions: both proteins bound angulin-3 60- to 140-fold more weakly than angulin-1, through different kinetic mechanisms. Ang1’s preference arose chiefly from 24-fold slower association, with dissociation contributing only 6-fold, whereas Ang2’s reflected changes in both rates, with 12-fold faster dissociation making the larger contribution (Supplementary Tables 1 and 2).

### *In silico* saturation mutagenesis maps determinants of angulin recognition

To identify positions where the angulin interfaces respond differently, we performed *in silico* saturation mutagenesis across the predicted interface (Figure 3A). We designate the Ib545–664 fragment Ang2 and number it from Ib545 as position 1; all residue numbers refer to Ang2 unless otherwise indicated (Ang2 position = Ib number − 544). Our previous molecular dynamics (MD) simulations resolved contact residues in Ang1 complexes with angulin-1 or angulin-3^23^, and we separately identified six Ang1 residues indispensable for angulin binding^13^. Of the MD contact residues, Ang1 Y643, corresponding to Ang2 Y99, was most critical for binding^23^.

**Figure 3.**
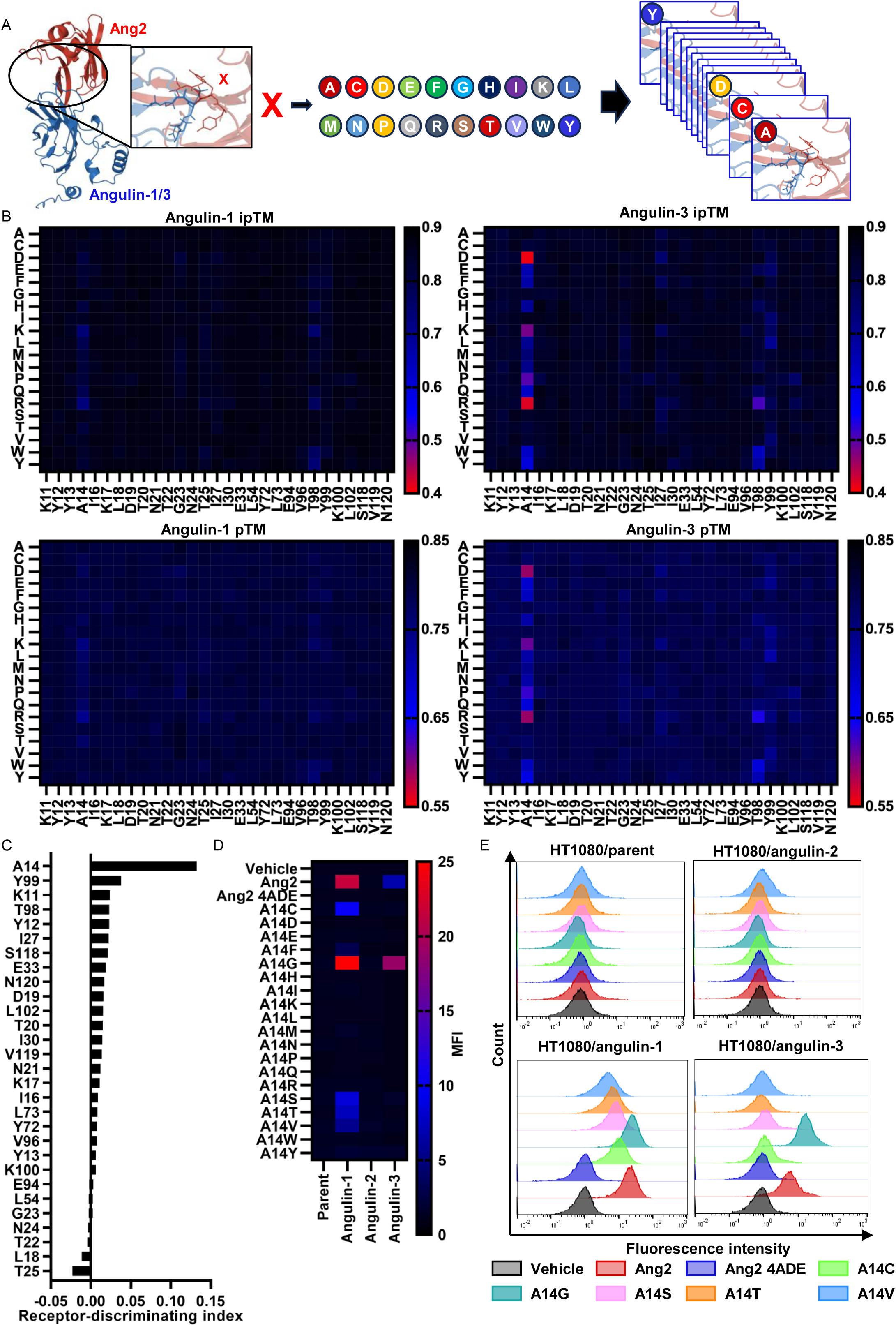
Ang2–angulin interface saturation substitution scan and A14 mutant selectivity. (A) *In silico* saturation substitution scan; Ang2 is red and angulin-1/-3 ECD blue. (B) Interface (ipTM, upper) and whole-complex confidence (pTM, lower) for Ang2–angulin-1 (left) and Ang2–angulin-3 (right). Columns denote Ang2 positions; rows denote substituting amino acids in one-letter code. Native-residue cells provide each column’s wild-type reference. Color scales: ipTM, 0.4–0.9; pTM, 0.55–0.85. A14 alone differs between receptors: angulin-3 ipTM falls to 0.41 (A14D), while angulin-1 ipTM remains ≥ 0.73. (C) Positions ranked by receptor-discriminating index. For each of 29 positions, ΔipTM equals wild-type ipTM minus mean ipTM across 19 substitutions, calculated separately per receptor. The index, ΔipTM(angulin-3) − ΔipTM(angulin-1), is positive when substitution costs more ipTM at angulin-3. A14 ranks first (0.133), 3.5-fold above Y99 (0.038). (D) Cell-surface binding of wild-type Ang2, interface-null Ang2 4ADE, and all 19 nonalanine A14 substitutions at 20 µg/mL to parental HT1080 cells and cells stably expressing FLAG-tagged angulin-1, angulin-2, or angulin-3. Bound protein was detected using an anti-His-tag antibody, biotinylated secondary antibody, and Alexa Fluor 647-conjugated streptavidin; DAPI-positive dead cells were excluded. Color denotes MFI of the stained population (scale, 0–25). All 21 proteins and vehicle were tested together on four lines in one experiment, representative of three independent experiments; text values are from this experiment. (E) Histograms from the experiment in (D) for seven indicated proteins and vehicle on each line.

We combined MD interface-contacting positions with the six essential residues to include positions implicated by either approach. Three essential residues (L54, E94, and V96 in Ang2 numbering) were not MD contacts and were added, yielding 29 positions (K11, Y12, Y13, A14, I16, K17, L18, D19, T20, N21, T22, G23, N24, T25, I27, I30, E33, L54, Y72, L73, E94, V96, T98, Y99, K100, L102, S118, V119, and N120). At each position, we introduced all remaining amino acids and re-modeled the resulting 551 mutants with angulin-1 and angulin-3 (Figure 3A). Absolute pTM and ipTM values are shown for both receptors (Figure 3B). For each position, we summarized substitution effects as ΔipTM, subtracting the mean interface confidence of its 19 substitutions from the native-residue value, and ranked positions by the receptor-discriminating index ΔipTM(angulin-3) − ΔipTM(angulin-1).

One position distinguished the receptors. With a discrimination index of 0.133, A14 ranked first, exceeding the next-ranked position, Y99 (0.038), by 3.5-fold (Figure 3C). A14 was also the most sensitive position for either receptor individually, and the next most responsive position was T98 — a position that has not been reported as functionally required (ΔipTM 0.045 for angulin-1 and 0.068 for angulin-3). However, T98 substitutions were nearly nondiscriminating (index 0.023, 5.7-fold below A14). Relative to the wild-type reference in the same scan (ipTM 0.87 for angulin-1 and 0.86 for angulin-3), substitution at A14 reduced the angulin-3 interface confidence by 0.184 on average across the 19 nonalanine substitutions and by up to 0.450 (ipTM 0.41 for A14D), whereas the same substitutions reduced the angulin-1 value by only 0.051 on average and 0.140 at most (ipTM 0.73 for A14R) (Figure 3B). We tested whether prediction stochasticity affected this asymmetry by repeating the complete A14 series with five independent seeds per complex (Supplementary Figure 1). Wild-type Ang2 gave an ipTM of 0.87 (range, 0.87–0.87) for the angulin-1 complex and 0.844 (range, 0.84–0.85) for the angulin-3 complex. Substitution effects exceeded seed-to-seed variation by an order of magnitude: the 19 nonalanine substitutions decreased the angulin-3 value by an average of 0.167 and a maximum of 0.414 (A14D, ipTM range 0.41–0.45).

We expressed and purified all 19 nonalanine A14 substitutions and wild-type Ang2, the alanine entry, and assayed cell-surface binding by FCM against HT1080 cells stably expressing angulin-1, angulin-2, or angulin-3 and against the parental line (Figure 3D, 3E). We report mean fluorescence intensities (MFI) of the stained population using the scale shown in Figure 3D; values for vehicle-treated cells ranged from 1.07 to 1.11 across the four lines. Wild-type Ang2 bound both angulin-1- and angulin-3-expressing cells strongly but gave no signal above background on angulin-2-expressing cells (1.20) or on the parental line (1.25); all proteins lacked detectable angulin-2 binding under these conditions. All four lines showed only background signal with the interface-null control Ang2 4ADE (1.03–1.14), confirming that the measured signal reflects the predicted interface. Substitution at A14 reorganized paralog preference. Five substitutions retained angulin-1 binding: A14G at 24.8, A14C at 10.3, A14S at 8.41, A14T at 7.28, and A14V at 5.49. Each exceeded its corresponding parental-cell signal by at least 3.6-fold. Angulin-3 binding was undetectable for 18 of the 19 substitutions (MFI ≤ 1.65 against a wild-type value of 6.76 and no more than 1.2-fold above the corresponding parental control). We therefore carried forward A14C, A14S, A14T, and A14V as a selective series: each retains 26–48% of wild-type angulin-1 binding while losing detectable angulin-3 binding. A14G alone retained angulin-3 binding; it bound angulin-1 at least as well as wild-type (24.8 vs. 21.5, 115%) but bound angulin-3 2.8-fold more strongly than wild-type (19.0 vs. 6.76), shifting preference in the direction opposite to the other four. These outcomes were not distinguished by prediction. The five substitutions that retained angulin-1 binding spanned an angulin-3 interface confidence of only 0.825–0.845, all within 0.04 of wild-type Ang2 (Figure 3B), so A14G was not separated from A14C, A14S, A14T, and A14V, which lost detectable angulin-3 binding.

### Kinetic analysis reveals the basis of angulin-1-selective binding

At the single protein concentration tested, FCM could neither distinguish lost angulin-3 affinity from weak binding below assay detection nor identify the reaction step affected by A14 substitutions. We therefore used SPR to measure affinity and binding kinetics against angulin-1 and angulin-3 (Supplementary Figure 2A, 2B) for wild-type Ang2 and the five mutants whose FCM profiles showed altered paralog preference: A14C, A14S, A14T, and A14V, which retained angulin-1 binding while losing detectable angulin-3 binding, and A14G, which retained angulin-1 binding while binding angulin-3 more strongly than wild-type.

A14G bound angulin-1 with near-wild-type affinity and showed no appreciable difference from wild-type Ang2 in either rate constant (Figure 4A, 4B, Table 1). A14C, A14S, A14T, and A14V had 5.0- to 13.9-fold weaker affinities. Each loss was driven almost entirely by faster dissociation: *k*_d_ increased 8.8- to 23.4-fold, whereas *k*_a_ increased at most 1.9-fold (Figure 4B, Table 1).

**Figure 4.**
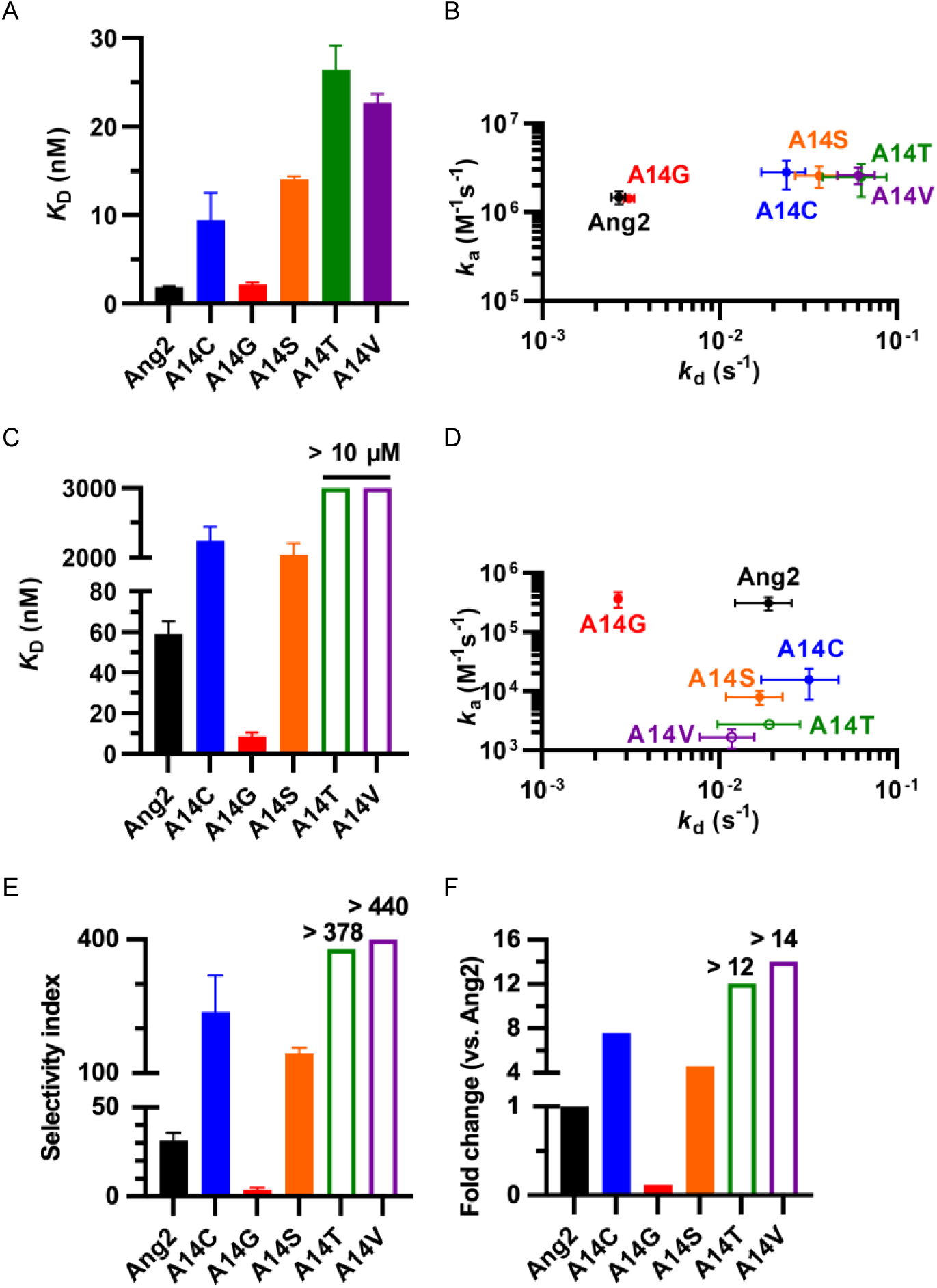
SPR resolves the kinetic basis of A14 mutant angulin-1 selectivity. His-tagged wild-type Ang2 or the indicated A14 mutant was captured on a Sensor Chip NTA, with human angulin ECD–rabbit Fc fusion proteins as analytes. (A, B) Angulin-1 equilibrium dissociation constant *K*_D_ (A) and association rate constant *k*_a_ vs. dissociation rate constant *k*_d_ (B). (C, D) Angulin-3 *K*_D_ (C) and *k*_a_ vs. *k*_d_ (D). Open bars in (C) mark A14T and A14V, whose affinity was not quantifiable up to 10 µM analyte; plotted values indicate *K*_D_ above that limit. Their rate constants were resolved in every replicate and are plotted as measurements in (D). Bars and symbols are arithmetic means ± SEM from three independent measurements. (E) Receptor selectivity, SI = *K*_D__angulin-3/*K*_D__angulin-1, calculated from (A) and (C). Bars are the ratio of the mean *K*_D_ values, with the error propagated. Open bars denote A14T/A14V lower limits calculated using *K*_D__angulin-3 > 10 µM, with annotations above bars. Note the segmented y-axis. (F) Data normalized to wild-type, SI(A14 mutant)/SI(Ang2); values above/below 1 indicate increased/decreased angulin-1 selectivity. Fold changes use mean indices. Open bars denote A14T/A14V lower limits calculated as in (E) and annotated above bars.

**Table 1.**
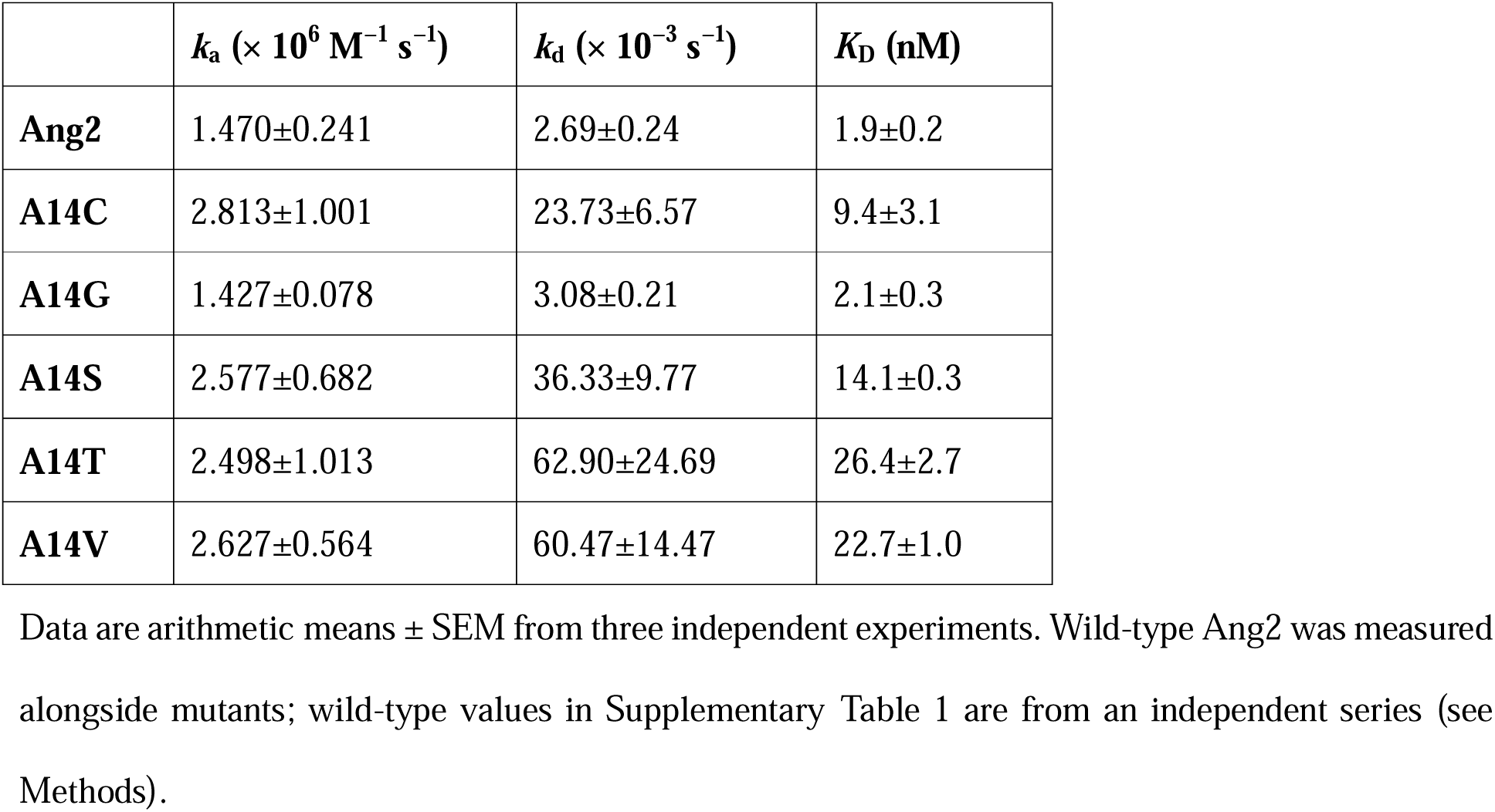
SPR-derived kinetic parameters for Ang2 and A14 mutants binding to human angulin-1.

|  | ***k*<sub>a</sub> (× 10<sup>6</sup> M<sup>−1</sup> s<sup>−1</sup>)** | ***k*<sub>d</sub> (× 10<sup>−3</sup> s<sup>−1</sup>)** | ***K*<sub>D</sub> (nM)** |
|---|---|---|---|
| **Ang2** | 1.470±0.241 | 2.69±0.24 | 1.9±0.2 |
| **A14C** | 2.813±1.001 | 23.73±6.57 | 9.4±3.1 |
| **A14G** | 1.427±0.078 | 3.08±0.21 | 2.1±0.3 |
| **A14S** | 2.577±0.682 | 36.33±9.77 | 14.1±0.3 |
| **A14T** | 2.498±1.013 | 62.90±24.69 | 26.4±2.7 |
| **A14V** | 2.627±0.564 | 60.47±14.47 | 22.7±1.0 |

Data are arithmetic means ± SEM from three independent experiments. Wild-type Ang2 was measured alongside mutants; wild-type values in Supplementary Table 1 are from an independent series (see Methods).

A14C and A14S bound angulin-3 with affinities roughly 35- to 38-fold weaker than that of wild-type Ang2. Here, the kinetic mechanism was reversed: *k*_a_ decreased 19.7-fold for A14C and 38.9-fold for A14S, while *k*_d_ remained within 1.7-fold of the wild-type value (Figure 4C, 4D, Table 2). Angulin-3 binding could not be quantified for A14T or A14V at concentrations up to 10 µM. Their fitted association rate constants were 113- and 187-fold below wild-type, whereas their fitted dissociation rate constants differed little from wild-type (Table 2). As the analyte series remained below saturation, we report affinity as a limit (*K*_D_ > 10 µM), but obtained resolved rate constants in every replicate — A14T’s association rate constant is the most precisely determined (SEM 3.2% of the mean, vs. 25.7% for wild-type Ang2) — and use these measured constants in the analyses below. A14G moved in the opposite direction, binding angulin-3 6.9-fold more tightly than the wild-type: its association rate constant was comparable to that of wild-type Ang2, whereas its dissociation rate constant fell 7-fold, accounting for the gain (Figure 4D, Table 2).

**Table 2.**
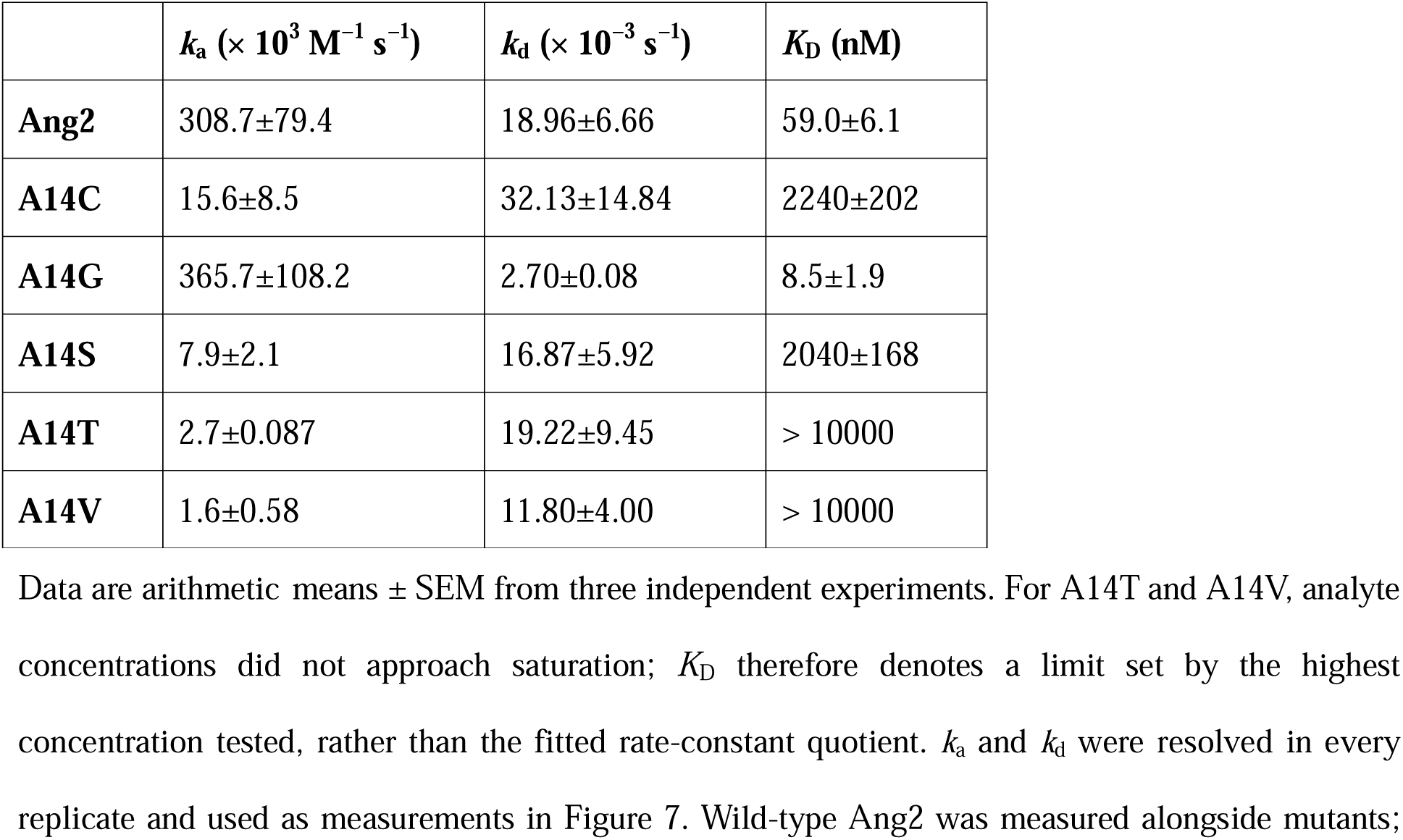
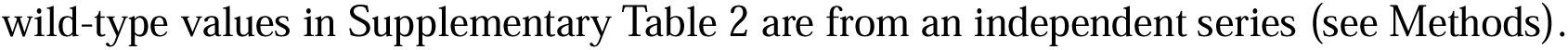
SPR-derived kinetic parameters for Ang2 and A14 mutants binding to human angulin-3.

We compared mutants using a selectivity index defined by the ratio of receptor *K*_D_ values obtained in the same experimental series (selectivity index, SI = *K*_D__angulin-3 / *K*_D__angulin-1; Tables 1 and 2, Figure 4E). Wild-type Ang2 preferred angulin-1 by 31-fold (SI = 31.4 ± 4.2). The four mutants lacking detectable angulin-3 binding in the FCM panel all increased SI: SI was 145 ± 12 for A14S and 238 ± 81 for A14C, and because A14T and A14V gave no quantifiable angulin-3 binding, their indices are lower limits (> 378 and > 440, respectively). Only A14G shifted preference in the opposite direction, largely eliminating the wild-type bias (SI = 3.9 ± 1.0) and yielding a fragment with nearly equal affinity for both receptors.

Normalizing indices to wild-type Ang2 places mutants on one selectivity scale (Figure 4F). Angulin-1 selectivity improved 4.6-fold for A14S and 7.6-fold for A14C, compared with more than 12-fold for A14T and 14-fold for A14V. A14G instead reduced selectivity 8.1-fold, to 0.12 of the wild-type value. A single substitution at position 14 can therefore switch selectivity in either direction, producing an angulin-1-restricted binder or a dual-receptor binder.

The kinetic basis of this switch differs between receptors. Faster dissociation accounted for almost all angulin-1 affinity loss in A14C, A14S, A14T, and A14V (Figure 4B), indicating that these side chains destabilize the complex but do not prevent it from forming. At the angulin-3 interface, association was chiefly affected (Figure 4D), consistent with the substituted side chain obstructing productive engagement. Selectivity is therefore gained by sacrificing angulin-3 engagement rather than by strengthening the angulin-1 interaction, which weakens in every selective mutant. A14T and A14V showed the greatest angulin-1 selectivity among the mutants characterized here, although their lower-limit indices prevent a ranking between them; their selectivity comes at the cost of absolute angulin-1 affinity; A14C and A14S offer intermediate trade-offs, retaining higher angulin-1 affinity with lower selectivity. By combining near-wild-type angulin-1 affinity with stronger angulin-3 binding, A14G instead favors applications requiring engagement of both receptors.

### Angulin-1-selective mutants modulate the Caco-2 barrier with distinct potencies

SPR measurements used purified receptor ECDs and do not establish whether mutants act on epithelia presenting angulins in their native context. We therefore used Caco-2 monolayers, with endogenous angulin-1 expression, to test the panel^23^. Monolayers grown on permeable supports were treated for 24 h with wild-type Ang2, interface-null Ang2 4ADE, or A14 mutants, and TER was measured before and after treatment; permeability to the 4-kDa tracer FD-4 was measured in the same monolayers to relate resistance to solute flux. At the initial test concentration of 50 µg/mL, the largest TER reduction was with A14G (72% of vehicle), followed by wild-type Ang2 (76%) and A14S (77%), then by A14T (84%) and A14V (89%), each differing significantly from vehicle; 4ADE (97%) and A14C (96%) were indistinguishable from vehicle (Figure 5A). FD-4 permeability increased in the same protein order, rising significantly above vehicle for A14G (2.4-fold), wild-type Ang2 (2.1-fold), A14S (1.9-fold), and A14T and A14V (1.7-fold each) (Figure 5B).

**Figure 5.**
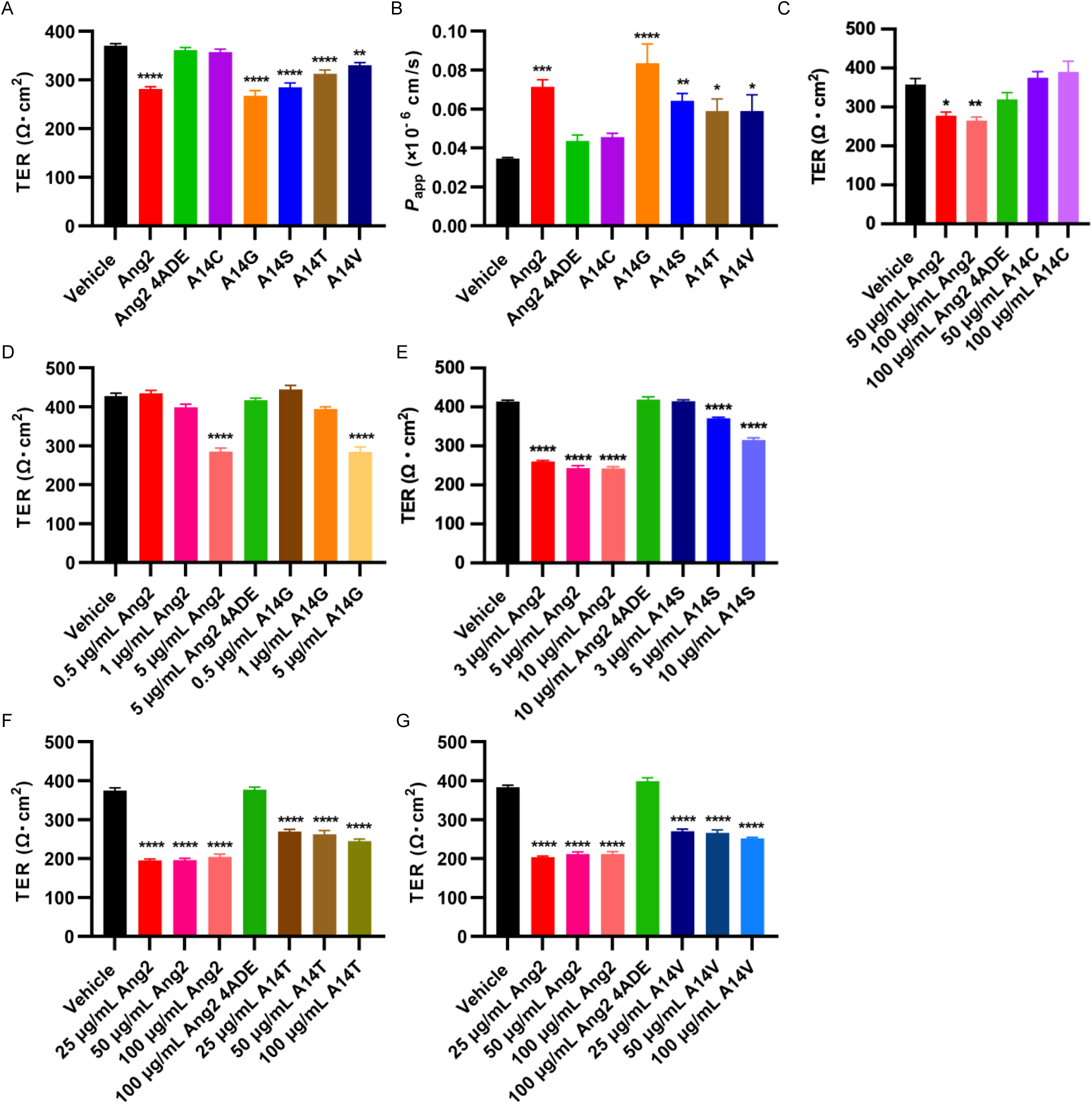
A14 mutant barrier modulation in Caco-2 monolayers is concentration-dependent and differs in potency. Mature Caco-2 monolayers on permeable supports were treated basally for 24 h with vehicle, wild-type Ang2, interface-null Ang2 4ADE, or the indicated A14 mutant. TER was measured before treatment and at 24 h. (A, B) Proteins at 50 µg/mL: TER at 24 h (A) and apparent permeability (*P*_app_) to 4-kDa FITC-dextran (FD-4) on the same monolayers (B). (C–G) Mutant concentration responses alongside wild-type Ang2 on the same plate; Ang2 4ADE at each series’ highest concentration is the negative control (green bars): A14C, 50 and 100 µg/mL (C); A14G, 0.5–5 µg/mL (D); A14S, 3–10 µg/mL (E); A14T, 25–100 µg/mL (F); A14V, 25–100 µg/mL (G). Bars are means ± SEM of replicate wells within a single experiment, each well a separate insert carrying its own monolayer; *n* = 5 wells per group in (A), (B), (E), (F), and (G), and *n* = 4 in (C) and (D). The experiment shown is representative of at least three independent repeats. Ordinary one-way ANOVA with Dunnett’s multiple-comparison test against vehicle: *, **, ***, \*\*\*\**P* < 0.05, 0.01, 0.001, 0.0001; unmarked groups did not differ from vehicle. Baseline TER did not differ across groups within any series (Supplementary Figure 3A–3F).

To resolve potency differences, mutants were titrated over individual ranges, with wild-type Ang2 assayed alongside on the same plate: A14G at 0.5–5 µg/mL, A14S at 3–10 µg/mL, A14T and A14V at 25–100 µg/mL, and A14C at 50–100 µg/mL. Within each series, treatment groups had comparable baseline TER (Supplementary Figures 3A–3F). A14C remained inactive across its full range, leaving TER unchanged at 50 and 100 µg/mL (105% and 109% of vehicle), while wild-type Ang2 on the same plate reduced it significantly to 78% and 74% (Figure 5C). The remaining four mutants reduced TER concentration-dependently (Figure 5D–5G), and their lowest active concentrations differed fivefold: 5 µg/mL for A14G and A14S, and 25 µg/mL for A14T and A14V. The most potent mutant, A14G, did not alter TER at 0.5 or 1 µg/mL (104% and 92% of vehicle) but significantly reduced it to 67% at 5 µg/mL, matching wild-type Ang2 tested alongside it at that concentration (67%) (Figure 5D). A14S left TER unchanged at 3 µg/mL (100%) and reduced it significantly to 90% at 5 µg/mL and to 76% at 10 µg/mL, where wild-type Ang2 had already plateaued at 59% (Figure 5E). The highest concentrations were required for A14T and A14V, which significantly reduced TER to 72% and 71% at 25 µg/mL and to 65% and 66% at 100 µg/mL, compared with 55% for wild-type Ang2 at 100 µg/mL; wild-type Ang2 had plateaued across 25–100 µg/mL in both series (Figure 5F, 5G). Ang2 4ADE left TER unchanged at each series’ highest concentration (98–104% of vehicle).

### Receptor rescue experiments establish angulin-1-dependent barrier modulation in EpH4 cells

Caco-2 monolayers express angulin-1 but neither angulin-2 nor angulin-3^23^, suggesting angulin-1-mediated activity. However, this assay provides neither a loss-of-function test nor information about the other paralogs. We therefore used an established EpH4 panel^11,13,23^ in which angulin-1 can be depleted by knockdown (KD) and angulin-2 or angulin-3 re-expressed in that background, isolating each receptor in its native epithelial tTJ context. The panel includes parental cells with endogenous angulin-1 expression (termed angulin-1-expressing EpH4 cells); angulin-1-KD cells; and angulin-1-KD cells with angulin-2 or angulin-3 re-expression (termed angulin-2-expressing and angulin-3-expressing EpH4 cells).

We treated monolayers with 100 µg/mL of wild-type Ang2, interface-null Ang2 4ADE, or one of four mutants: A14G, A14S, A14T, and A14V. Baseline TER did not differ across treatment groups within any series (Supplementary Figure 4A–4D). Wild-type Ang2 and all four mutants significantly reduced TER in angulin-1-expressing EpH4 cells: wild-type Ang2 to 55% of vehicle, A14G to 53%, A14S to 66%, A14T to 67%, and A14V to 55%, whereas 4ADE had no significant effect (Figure 6A). Receptor dependence was clear. TER remained at 98–108% of vehicle in angulin-1-KD EpH4 cells, with no treatment producing a change (Figure 6B). Thus, none perturbs the barrier in the absence of a cognate angulin, and the reductions seen in the other lines reflect receptor engagement rather than nonspecific monolayer effects. Likewise, no treatment significantly affected TER in angulin-2-expressing EpH4 cells, where values remained at 99%–110% of vehicle (Figure 6C), in agreement with the lack of angulin-2 binding detected by FCM.

**Figure 6.**
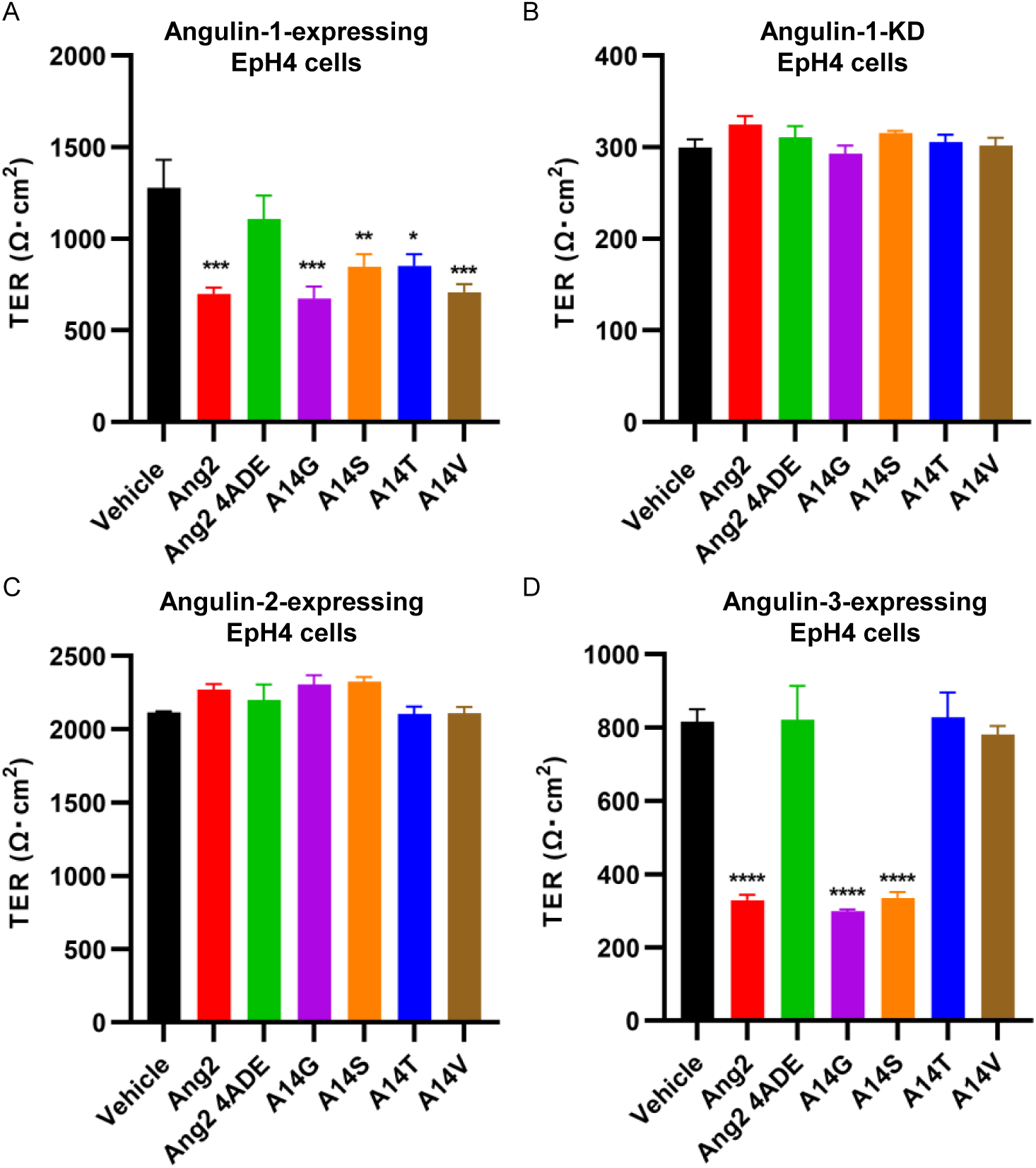
A14 mutant barrier modulation requires cognate angulin and differs most in angulin-3-expressing monolayers. EpH4 monolayers on permeable supports were treated basally for 24 h with vehicle, wild-type Ang2, interface-null Ang2 4ADE, or A14G, A14S, A14T, or A14V at 100 µg/mL. TER was measured before treatment and at 24 h; 24-h values are plotted. (A) Parental EpH4 cells expressing endogenous angulin-1. Wild-type Ang2 and all four mutants significantly reduced TER. (B) Angulin-1-KD EpH4 cells. No treatment altered TER, indicating dependence on cognate angulin. (C) Angulin-2-expressing EpH4 cells (angulin-1-KD cells re-expressing FLAG-tagged angulin-2). No treatment differed from vehicle. (D) Angulin-3-expressing EpH4 cells (angulin-1-KD cells re-expressing FLAG-tagged angulin-3). Wild-type Ang2, A14G, and A14S significantly reduced TER; A14T, A14V, and Ang2 4ADE did not. Bars are means ± SEM of replicate wells within a single experiment, each well a separate insert carrying its own monolayer; *n* = 5 per group. The experiment shown is representative of at least three independent repeats. Ordinary one-way ANOVA with Dunnett’s multiple-comparison test against vehicle: *, **, ***, \*\*\*\**P* < 0.05, 0.01, 0.001, 0.0001; unmarked groups did not differ from vehicle. Baseline TER did not differ across groups within any series (Supplementary Figure 4A–4D).

The clearest distinction between mutants emerged in angulin-3-expressing EpH4 cells. Wild-type Ang2 significantly reduced TER to 40% of vehicle, and A14G and A14S were at least as active (37% and 41%). In contrast, A14T, A14V, and Ang2 4ADE were inactive, leaving TER at 101%, 96%, and 101%, respectively (Figure 6D).

These results establish a functional basis for the SPR selectivity. A14T and A14V, whose angulin-3 affinities were below quantification, modulated the barrier through angulin-1 alone; A14G and A14S retained measurable angulin-3 binding and acted through both receptors. Thus, A14T and A14V confer functional angulin-1 selectivity on Ang2. A14G is a more potent dual-receptor modulator, retaining near-wild-type angulin-1 activity while binding angulin-3 more tightly and acting more strongly on angulin-3-expressing monolayers than wild-type.

### Side-chain bulk at position 14 determines which kinetic step governs angulin selectivity

The kinetic measurements distinguish the reaction step affected at each receptor, but they do not identify the responsible side-chain property. The six residues examined differ systematically in their number of side-chain heavy atoms beyond Cβ: zero for glycine and wild-type alanine, one for cysteine and serine, and two for threonine and valine. Because these side chains do not extend beyond the γ position, the count also represents the number of γ atoms, denoted γ in the figures. Ranked on this count, the two rate constants governing selectivity were ordered monotonically — the dissociation rate constant at angulin-1 (Spearman ρ = +0.956) and the association rate constant at angulin-3 (ρ = −0.956), each with exact permutation *p* = 0.0222 (Figure 7A, 7B). No such ordering was evident for the two constants that do not govern selectivity: the association rate constant at angulin-1 and the dissociation rate constant at angulin-3 (Supplementary Figure 5A, 5B). Grouping proteins by the presence of any atom beyond Cβ gives 74.8-fold slower association at angulin-3 and 13.4-fold faster dissociation at angulin-1.

**Figure 7.**
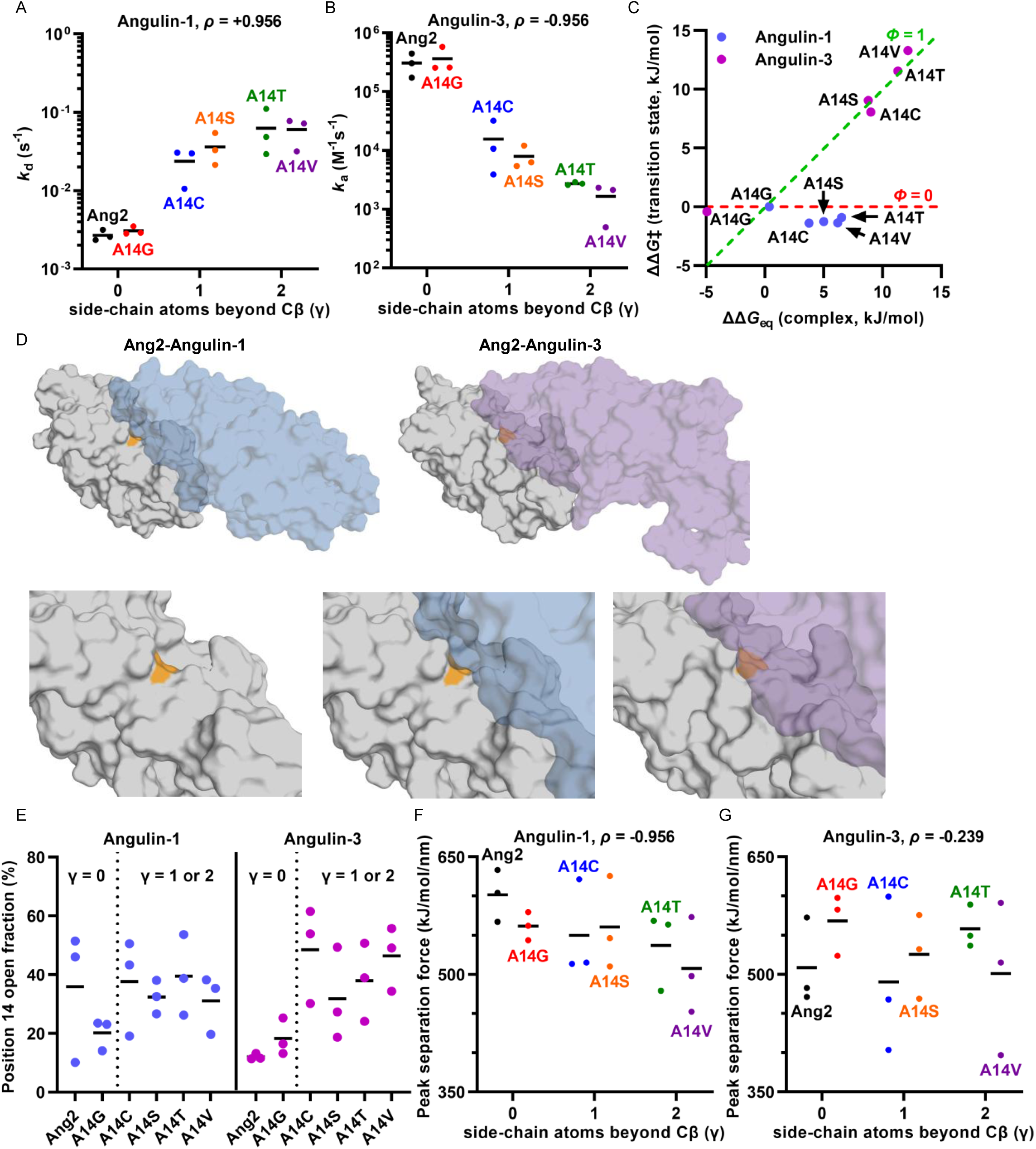
Side-chain atom count beyond Cβ at position 14 determines the kinetic step affected at each receptor. (A, B) Selectivity-determining rate constants vs. side-chain heavy-atom count beyond Cβ (γ): angulin-1 dissociation (A) and angulin-3 association (B). Three independent SPR determinations per protein; horizontal lines are arithmetic means of the three determinations; ρ denotes Spearman’s correlation across six proteins; ρ was calculated on the geometric means and is identical for the arithmetic ones. Other rate constants appear in Supplementary Figures 5A and 5B. (C) Brønsted plot of transition-state penalty ΔΔ*G*‡ vs. overall penalty ΔΔ*G*_eq_; both quantities are named in words on the axes. Blue, angulin-1; magenta, angulin-3. The diagonal (Φ = 1) indicates the whole penalty at the transition state; the horizontal (Φ = 0) indicates none. Points are formed from the mean of ln *k*_a_ and the mean of ln(*k*_a_/*k*_d_) across the three replicates. (D) Representative wild-type MD conformations in surface representation: Ang2, gray; angulin-1, blue; angulin-3, purple; position 14, orange. Top, common orientation after superposing Cα atoms of 18 interface residues; bottom, close views of position 14 in Ang2 alone and with each receptor. Angulin-3 covers position 14, whereas position 14 remains at the angulin-1 interface rim. (E) Fraction of MD frames with any position 14 solvent exposure at angulin-1 (left) or angulin-3 (right). Three independent 100-ns replicas per system; bars are arithmetic means across replicas; the dotted line separates γ = 0 from γ = 1 or 2. (F, G) Peak separation force for angulin-1 (F) and angulin-3 (G), averaged across six starting conformations per replica; bars are arithmetic means across replicas, and ρ is as in (A, B).

The two rate constants locate each penalty along the reaction coordinate. For each substitution, we calculated the transition-state penalty ΔΔ*G*‡ from the association rate constant and the overall penalty ΔΔ*G*_eq_ from the equilibrium constant assembled as *k*_a_/*k*_d_ within each replicate; all quantities entering Φ therefore derive from measured rate constants, excluding the angulin-3 affinity limits for A14T and A14V. The ratio of these penalties, Φ = ΔΔ*G*‡/ΔΔ*G*_eq_, quantifies how much of the final penalty is present at the transition state. At angulin-3, Φ ≈ 1 for A14C, A14S, A14T, and A14V: the penalty is complete by the transition state, with no further penalty upon complex formation (Figure 7C). The same substitutions gave Φ ≈ 0 at angulin-1, placing the penalty entirely in the bound complex rather than at the transition state (Figure 7C). Because the free state of Ang2 is shared between the two receptors, a substitution that impaired the free protein would slow association at both. Association at angulin-1 is unaffected, both in Φ and in its lack of any atom-count ordering (Supplementary Figure 5A), so the angulin-3 association penalty arises in the encounter with that receptor rather than in free Ang2. A14G, the only substitution that improves affinity, does so at angulin-3 (ΔΔ*G*_eq_ = −4.96 kJ/mol) with Φ = 0.08, indicating essentially no change at the transition state, consistent with a gain arising entirely from complex lifetime. The two receptors are therefore discriminated by when the substitution’s cost is incurred rather than its magnitude.

*Φ* ≈ 1 at angulin-3 means that the full cost of substitution is expressed by the barrier and none is added on complex formation. This supports an inference about energetics: the feature responsible for the cost is already in effect when the barrier is crossed. It does not specify the structure of the transition state, since Φ cannot distinguish a complex-like environment at position 14 from any other arrangement of equal cost. We used bound-state simulations to characterize the environment that position 14 occupies once bound; these simulations sample equilibrium configurations of the complex, not the barrier, and therefore describe that environment rather than quantify the penalty. Three independent 100-ns simulation replicas were run for each of the 12 complexes: wild-type Ang2 and five substitutions paired with each receptor. Position 14 lies within the contact in both complexes and is enclosed by the receptor: its median per-frame exposure is zero in all twelve systems. Enclosure differs between receptors. Trajectory-averaged exposure at wild-type position 14 is 20.3% of its free-state value at angulin-1, compared with only 4.6% at angulin-3. Representative conformations from the simulations show this asymmetry: at angulin-3, the receptor surface closes over position 14, whereas the residue remains at the rim of the interface at angulin-1 (Figure 7D).

Exposure of position 14 was intermittent rather than graded: the residue is completely enclosed for 51%–88% of the trajectory and opens to 38%–66% of its free-state exposure for the remainder, so opening frequency distinguishes proteins better than a single exposure value. At angulin-3, the two proteins lacking an atom beyond Cβ opened in 15.2% of frames and the other four in 41.2%, with no overlap between the protein means (a gap of 2.2 SEM) (Figure 7E). At angulin-1, the corresponding values were 28.0% and 35.2%, respectively, and the groups overlapped, so this receptor does not distinguish the groups — the contrast is confined to angulin-3. Wild-type provides the baseline for the added atom’s effect: alanine opened in 12% of frames at angulin-3 but in 36% at angulin-1, indicating that the tighter angulin-3 enclosure leaves little room for an atom beyond Cβ. A similar contrast emerges when time-averaged position 14 exposure is expressed relative to its free-state value (Supplementary Figure 5C). The effect does not run through the dominant contact of the interface: Y99, the residue most critical for angulin binding in the parent molecule^23^, was in contact in every frame of all twelve systems without on–off transitions, and counting contact atom pairs showed more contacts in proteins losing angulin-3 association than in wild-type.

If an atom beyond Cβ destabilizes the bound angulin-1 complex, separating that complex should require less force. Each system was pulled apart from six conformations per replica, recording peak force. Peak force at angulin-1 declined monotonically as atom count increased (ρ = −0.956), in agreement with faster dissociation measured by SPR; no such decline occurred at angulin-3 (ρ = −0.239) (Figure 7F, 7G), as expected if the cost of the substitution at this receptor is already paid before the complex forms.

Together, the number of side-chain heavy atoms beyond Cβ at position 14 — taken here as an operational measure of side-chain bulk — is the property affecting the two receptors at different reaction steps. At angulin-3, the atom imposes its cost on a step completed before the complex forms; at angulin-1, it is accommodated but destabilizes the complex.

## Discussion

This study sought a paralog-discriminating modulator of tTJs for an interaction lacking an experimental structure. We used AF3 to nominate rather than determine the interface, treating every prediction as a hypothesis for protein-level testing: the scan defined a minimal functional scaffold, Ang2, and identified the single position where the two angulin interfaces diverge.

The performance of the scan informs the use of interface prediction in design. Ranking the scanned positions by their differential predicted effects on the receptors placed position 14 first, 3.5-fold above the next position, and that ranking was correct. Within that position, however, the prediction had no resolving power: the five substitutions supporting the selectivity conclusion are indistinguishable in predicted interface confidence yet yield three experimental outcomes. The scan also failed to flag five of the six residues that alanine scanning identified as required for angulin binding^13^, and interface confidence moved in lockstep with whole-complex confidence across the series. It therefore measured complex prediction confidence rather than binding energetics, consistent with the finding that AlphaFold confidence does not track the effects of point substitutions on binding^19^. The two failures are of different kinds. The index that selected position 14 was differential — the difference between two receptors for the same ligand sequence, which cancels whatever the confidence shares across them. The index that failed was absolute — whether a given residue is required for binding at all. A differential ranking can therefore be informative while the underlying absolute values are not, and only the differential form was used here.

Prediction succeeded despite the fact that no structure of an angubindin–angulin complex exists and bacterial toxin and mammalian receptor lack a shared evolutionary history permitting inter-chain covariation. The properties associated with accurate nonantibody heterodimer prediction — buried interface area, interface rigidity, and representation of the ligand’s family in the training set — are present in this interface^24–28^, and its confidence of 0.86 exceeds the empirical acceptance threshold of 0.70 for AF3^25^. Confidence is not self-validating^29^, but here it distinguished three homologous receptor ectodomains, collapsing to 0.19 for angulin-2 and correctly predicting that angulin-2 is not engaged. The prediction supported the fold, domain boundary, and interface location, the levels at which we used it.

SPR identified different kinetic steps underlying receptor discrimination: an atom beyond Cβ at position 14 determines which step carries the cost. At angulin-3, the penalty falls entirely on association, which slows 74.8-fold, costing 10.7 kJ/mol at the barrier; at angulin-1, the same atom leaves association unchanged and accelerates dissociation 13.4-fold, costing 6.4 kJ/mol only in the bound complex. A penalty confined to association, with the lifetime of the complex preserved, indicates a substitution impeding the encounter complex or the transition state rather than destabilizing the bound state^30,31^, and Φ locates that penalty without requiring a structure of either transition state. Φ ≈ 1 at angulin-3 constrains the energetics of that step, not its structure: the feature responsible for the cost is already in effect when the barrier is crossed, but Φ cannot distinguish a complex-like environment at position 14 from any other arrangement of equal cost. The bound-state simulations therefore supply a candidate for that feature rather than a view of the barrier. In the complex, the receptor surface closes over position 14, which retains 4.6% of its free-state exposure against 20.3% at angulin-1, and the enclosure opens in 15.2% of frames without an atom beyond Cβ and 41.2% with one, favoring an ensemble of open and closed states over a single structure^32^. A pocket this restrictive once bound is a plausible origin for a cost incurred on approach, but the simulations do not establish that it is. This comparison does not distinguish proteins at angulin-1, where the distinction is mechanical: the force needed to separate the complex falls monotonically with the atom count, as the measured dissociation predicts. The effect does not run through the dominant contact of the interface; Y99 remains engaged in every frame of every system. Glycine is the reciprocal test, and the two directions are not symmetric: removing the Cβ slows dissociation 7.0-fold without accelerating association, raising angulin-3 affinity 6.9-fold with the barrier unchanged (Φ = 0.08). Position 14 therefore determines the kinetic step affected through two features — the height of the angulin-3 association barrier and the depth of the angulin-1 well — providing two independent settings for selectivity. The distinction is practical because affinity gains through complex lifetime and association have different implications for the duration of a barrier-opening effect^33^.

Binding selectivity translated into functional selectivity. In the EpH4 panel, A14T and A14V reduced resistance in angulin-1-expressing monolayers to 66.6% and 55.4% of vehicle while leaving angulin-3-expressing monolayers at 101.4% and 95.7%, establishing functional paralog selectivity; A14G and A14S remained active on both receptors, consistent with retained angulin-3 binding. Because angulin-3 is presented by an overexpressing rescue line, the failure of A14T and A14V to act there understates their selectivity, whereas the activity of A14G and A14S may be enhanced by overexpression; the selectivity shown here is therefore established at rescue-line rather than endogenous receptor densities, requiring matched densities or an epithelium expressing angulin-3 endogenously. Affinity does not explain one mutant. A14C binds angulin-1 more tightly than any other selective mutant yet left Caco-2 resistance unchanged at 100 µg/mL, 20 times the active A14S concentration. Occupancy cannot account for that, and because interface-null 4ADE is also inactive, engagement of angulin-1 is necessary but not sufficient for barrier modulation. The basis for this remains unresolved, and the A14C thiol is a candidate: Ang2 has no native cysteine, so the substitution introduces a single unpaired thiol, susceptible to oxidation or mixed-disulfide formation. Neither binding assay reproduces functional-assay conditions: SPR exposed the captured protein for a short period on a surface renewed every cycle, and FCM staining was done on ice for over 30 min, whereas the functional readout followed monolayers for 24 h at 37 °C in serum-containing medium.

Paralog engineering of TJ-binding bacterial proteins has precedent in the claudin field, where C-CPE has yielded binders with altered claudin preference^15,34^, and claudin-5-directed binders have been applied to the blood–brain barrier^35,36^. In the angulin family, the precedent is combinatorial diversification of the shared binding residues of Ang1 itself^23^, which yielded an angulin-1-selective variant without nominating a position in advance. This family is of interest because the three paralogs have distinct tissue roles: angulin-1 organizes tTJs at the blood–brain barrier^7,12^, angulin-2 is required for hearing^8^ and mediates fat-stimulated cholecystokinin secretion^37^, and angulin-3 participates in hepatic lipid homeostasis^38^. A modulator restricted to one paralog would therefore be expected to have a different tissue footprint from the parent molecule, which engages two of the three; testing this *in vivo* remains beyond this study.

Several limitations qualify these conclusions. The mutated interface is predicted: the functional and kinetic data support a shared interface that includes position 14 and has receptor-dependent effects, but they do not validate the atomic details of the models, and confidence values are used only as rankings. The absence of angulin-2 engagement and angulin-3 binding for the selective variants are statements about the conditions tested and do not exclude weak interactions. Six proteins constrain the kinetic series: the contrast between the presence and absence of an atom beyond Cβ is established, whereas the monotonic ordering by atom count reaches an exact permutation *p* of 0.0222, the smallest attainable value for six proteins ranked on a three-valued predictor. The count also incompletely describes the side chain: a size series would distinguish its effect from side-chain volume. Association at angulin-1 is not entirely flat either, rising 1.65-fold when an atom beyond Cβ is present, an effect five- to nine-fold smaller in energy than the two that carry the selectivity and consistent with slightly negative Φ at that receptor. No trajectory reaches the association barrier: the simulations sample equilibrium configurations of the bound complexes, so the enclosure they describe stands as an inference about that barrier, supported by Φ, rather than an observation of it. The simulations were run as three 100-ns replicas per system, limiting interpretation of null results, including absent discrimination at angulin-1. They were analyzed on protein coordinates, leaving water-mediated contributions unexamined. Separation at 0.5 nm/ns is far from equilibrium, so only peak-force ranks are interpreted. Finally, all functional data come from cultured monolayers.

These results support a division of labor between prediction and experiment for structurally unresolved interfaces: use the prediction to identify where homologous interfaces diverge, and use protein-level binding and function to determine the substitution. This logic should apply wherever a receptor family must be discriminated and no structure of the relevant complex exists. It is complementary to *de novo* binder design^20,21^, which requires a target structure. Immediate questions are whether the affinity lost at angulin-1 can be recovered by maturation elsewhere in the footprint without restoring angulin-3 engagement, and whether a paralog-biased modulator opens the blood–brain barrier with a narrower peripheral footprint than Ang1^11,12^. Because the three angulins occupy different tissues, a modulator that engages angulin-1 while sparing angulin-3 is the form in which tTJ opening becomes targetable rather than generalized. Ang2 provides a compact scaffold for this purpose, and A14T and A14V show that a single substitution is sufficient to obtain one: the same position sets both the receptor preference and the kinetic step that carries it. Because discrimination against angulin-3 and the lifetime of the angulin-1 complex reside in different rate constants, selectivity and the duration of the effect can be engineered separately.

## Materials and Methods

### Structure prediction using AlphaFold3

All structure predictions were generated with AlphaFold3 (AF3) via the AlphaFold Server (https://alphafoldserver.com)^18^. The *Clostridium perfringens* iota-toxin receptor-binding component Ib (NCBI Reference Sequence WP_003463384.1) served as the ligand. Angubindin-1 numbering (Ang1; Ib421–664) follows mature, proteolytically activated Ib, as established previously^13^, and differs by 211 residues from the 875-residue precursor containing an N-terminal signal peptide and propeptide. Thus, Ang1 corresponds to mature Ib residues 421–664 and precursor residues 632–875. Angubindin-2 (Ang2) comprises Ib545–664 (120 aa) and is numbered 1–120; Ang2 residue *n* corresponds to Ib residue *n* + 544 (e.g., A14 = A558, N120 = N664). Only human angulin ECDs were submitted because transmembrane (TM) and cytoplasmic regions do not participate in angubindin binding and introduce large disordered segments that depress global confidence metrics: angulin-1 (NP_991403.2, residues 42–211; signal peptide omitted, terminating immediately before the TM segment), angulin-2 (NP_001186728.1, residues 24–167; likewise), and angulin-3 (NP_955383.1, residues 21–186; likewise). Each job used the default single seed, returning five diffusion samples. All samples and confidence files were downloaded and retained as the job ZIP archive. Predictions were assessed using the server’s four confidence measures: per-atom pLDDT, predicted aligned error (PAE), and global pTM and ipTM. Models were ordered by ranking score, which also penalizes steric clashes and spurious helicity in disordered regions; this score was used only within jobs, never between jobs.

Twenty-nine Ang2 positions were selected for *in silico* saturation mutagenesis using two criteria. First, alanine scanning of Ang1 identified six residues essential for angulin binding: L18, L54, E94, V96, Y99, and K100 in Ang2, corresponding to L562, L598, E638, V640, Y643, and K644 in Ang1^13^. Second, our previous MD simulations resolved predicted Ang1 interfaces with angulin-1 and angulin-3^23^. The overlapping criteria yielded 29 unique positions: K11, Y12, Y13, A14, I16, K17, L18, D19, T20, N21, T22, G23, N24, T25, I27, I30, E33, L54, Y72, L73, E94, V96, T98, Y99, K100, L102, S118, V119, and N120.

Mutant sequences were generated by editing the Ang2 input sequence without templates, structural restraints, or other modeling constraints. Each mutant was co-folded separately with angulin-1 and angulin-3 ECDs using the wild-type Ang2 settings. The top-ranked sample’s pTM and ipTM values were extracted from each job’s summary confidences file. At each position, ΔipTM was calculated as native-residue ipTM minus mean ipTM across its 19 substitutions, separately for each receptor; positions were ranked by ΔipTM(angulin-3) − ΔipTM(angulin-1). To test robustness to prediction stochasticity, the complete A14 series was repeated with five independent seeds per complex (Supplementary Figure 1).

### Cell culture

HT1080 cells and HT1080 cells overexpressing FLAG-tagged angulin-1 (LSR), angulin-2 (ILDR1), or angulin-3 (ILDR2)^39^ were maintained in Dulbecco’s modified Eagle’s medium (DMEM) with 1% penicillin–streptomycin and 10% fetal bovine serum (FBS) at 37 °C under 5% CO_2_. Caco-2 cells were maintained in minimum essential medium (MEM) with 1% penicillin–streptomycin, 10% FBS, and 1% MEM Nonessential Amino Acids Solution (Nacalai Tesque, Kyoto, Japan) under the same conditions. Mouse mammary epithelial EpH4 cells (angulin-1-expressing), angulin-1-knockdown EpH4 cells (angulin-1-KD), and angulin-1-KD cells overexpressing FLAG-tagged angulin-2 or angulin-3 (angulin-2-expressing or angulin-3-expressing, respectively) were kindly provided by Dr. Mikio Furuse (Kobe University, Japan)^5^. All EpH4 cells were maintained in DMEM with 1% penicillin–streptomycin and 10% FBS at 37 °C under 5% CO_2_.

### Purification of recombinant Ang1 and Ang2

Recombinant Ang1 and Ang1 4ADE were expressed in *E. coli* as N-terminally His-tagged proteins as described previously^23^. The pET-16b-Ang2 plasmid was assembled from two PCR fragments using In-Fusion cloning. The vector fragment was amplified from pET-16b (Merck KGaA, Darmstadt, Germany) using forward primer TAAGGATCCGGCTGCTAACAAAGC and reverse primer CATATGACGACCTTCGATATGG. The Ang2 fragment was amplified from pET-16b-Ang1^23^ using forward primer GAAGGTCGTCATATGAGTGACCAAGAAATACTAGATGC and reverse primer GCAGCCGGATCCTTAATTAACACTAAGCACTAATAACTC. Products were resolved by agarose gel electrophoresis, excised, purified, and joined using an In-Fusion cloning kit (Takara Bio, Shiga, Japan). The pET-16b-Ang2 4ADE plasmid was constructed similarly from pET-16b-Ang1 4ADE using the same primers. Other Ang2 mutant expression plasmids were generated by PCR with primers in Supplementary Table 3. All plasmids were transformed into *E. coli* BL21(DE3) (Merck KGaA).

Transformed bacteria were cultured in Terrific Broth containing ampicillin (25 µg/mL) at 37 °C. At an optical density of 0.4–0.6 at 600 nm, expression was induced with 1 mM isopropyl-β-D-thiogalactopyranoside. Cultures were incubated for another 20 h at 25 °C, and cells were harvested by centrifugation. Cells were lysed by sonication in buffer A (25 mM Tris-HCl [pH 8.0], 400 mM NaCl, 5 mM MgCl_2_, 10% glycerol), and lysates were clarified by centrifugation at 4 °C.

Recombinant proteins were purified from clarified supernatants under native conditions using HisPur Cobalt Resin (Thermo Fisher Scientific). After loading, bound proteins were eluted with buffer A containing 100 mM imidazole.

Pooled fractions were buffer-exchanged into phosphate-buffered saline (PBS) using Amicon Ultra-4 centrifugal filters (10-kDa MWCO; Merck Millipore) to remove imidazole. Purified proteins were aliquoted and stored at −80 °C. Concentrations were determined by absorbance at 280 nm, and purity was assessed by SDS-PAGE.

### FCM analysis

Recombinant protein binding was evaluated by FCM. Cells were harvested and incubated in FACS buffer (PBS, 1% FBS, 2 mM EDTA) on ice for 30 min. His-tagged proteins were added, and cells were incubated on ice for 30 min. Final protein concentrations were 20 µg/mL for Ang2–mutant comparisons and 1 µM for Ang1–Ang2 comparisons. After washing in FACS buffer, cells were incubated with an anti-His-tag monoclonal antibody (clone H-3, cat. sc8036, Santa Cruz Biotechnology, Dallas, TX, USA) on ice for 30 min. Cells were washed again and incubated with biotin-SP (long spacer)-conjugated AffiniPure donkey anti-mouse IgG (H+L) polyclonal antibody (cat. 715-065-151, Jackson ImmunoResearch Laboratories, West Grove, PA, USA) on ice for 30 min. After another wash, Alexa Fluor 647-conjugated streptavidin (BioLegend, San Diego, CA, USA) and DAPI (Dojindo Laboratories, Kumamoto, Japan) were added; DAPI-positive dead cells were excluded from analysis. Cells were incubated on ice for 20 min protected from light. After a final wash, fluorescence was measured using a MACSQuant X flow cytometer (Miltenyi Biotec, Bergisch Gladbach, Germany) and analyzed using FlowJo v10.8.1 (BD Biosciences, Franklin Lakes, NJ).

### Measurement of transepithelial electrical resistance (TER) and permeability assays

Cell culture inserts (Falcon Permeable Support for 24-well Plate with 0.4-µm Transparent PET Membrane, Corning) were placed in 24-well plates. EpH4 and Caco-2 cells were seeded apically at 1 × 10^4^ and 6 × 10^4^ cells per insert, respectively, in 500 µL of medium, with 800 µL added basally. Cells were cultured at 37 °C under 5% CO_2_; TER was monitored every 2–3 days using a voltohmmeter (Millicell ERS 3.0, Merck Millipore). Caco-2 cells and angulin-2- or angulin-3-expressing EpH4 cells were assayed on day 10; angulin-1-expressing and angulin-1-KD EpH4 cells were assayed on day 7.

To evaluate tTJ modulation, recombinant proteins were added basally at 0.5–100 µg/mL; control wells received an equivalent PBS volume. Subsequently, 500 µL of medium was added apically. TER measured immediately before protein addition defined the 0-h value, with subsequent measurements at 3, 6, 9, and 24 h.

After 24 h of treatment, 200 µL of fluorescein isothiocyanate-labeled dextran (FD-4; 4 kDa, 2.5 mg/mL) in phenol red-free medium was added apically to assess paracellular permeability. After 4 h, basal medium was sampled and fluorescence measured at excitation/emission wavelengths of 485/535 nm using a fluorescence spectrophotometer (TriStar LB941, Berthold Technologies, Bad Wildbad, Germany). FD-4 concentrations were calculated from a serial-dilution standard curve.

### Purification of recombinant angulin-1 and angulin-3

Human angulin-1 and angulin-3 ECD–rabbit Fc fusion proteins were expressed in Expi293F cells by transient transfection of the corresponding pcDNA3.4 plasmids using ExpiFectamine 293 (Thermo Fisher Scientific), as described previously^23^. ECD expression plasmids were prepared as described previously^23^. Supernatants were collected 6 days after transfection, and secreted proteins were purified by Protein A affinity chromatography. Protein A beads were incubated with supernatants at 4 °C for at least 3 h, washed with PBS, and eluted with 0.1 M glycine-HCl (pH 3.0). Eluates were immediately neutralized with one-tenth volume of 1 M Tris-HCl (pH 9.0). Proteins were buffer-exchanged using Amicon Ultra-4 centrifugal filters (10-kDa MWCO; Merck Millipore) into HBS-P+ for angulin-1 or HBS-P+ with an additional 350 mM NaCl (500 mM final) for angulin-3. Concentrations and purity were determined by absorbance at 280 nm and SDS-PAGE, respectively.

### Surface plasmon resonance (SPR)

Binding kinetics of recombinant Ang1, Ang2, and Ang2 mutants to human angulin-1 and angulin-3 were analyzed by SPR using a Biacore 1S+ system and Series S Sensor Chip NTA (Cytiva, Uppsala, Sweden). His-tagged proteins purified from *E. coli* as above were captured using 0.5 mM NiCl_2_ (60-s injection, 5 μL/min); the reference flow cell lacked NiCl_2_ and ligand. Human angulin-1 and angulin-3 ECD–rabbit Fc fusion proteins purified from Expi293F cells served as analytes. Ligands and analytes were buffer-exchanged into the appropriate running buffer and centrifuged or filtered to remove aggregates.

Multicycle SPR measurements were performed at 25 °C and 30 μL/min. For human angulin-1 ECD-Fc, the running buffer was HBS-P+ (10 mM HEPES, 150 mM NaCl, 0.05% Surfactant P20, pH 7.4). Analyte serial dilutions of 1.23–300 nM were injected for 120 s, followed by 600 s of dissociation. For angulin-3 ECD-Fc, HBS-P+ contained an additional 350 mM NaCl (500 mM final) and 0.5 mg/mL BSA to suppress nonspecific surface binding. Analyte serial dilutions of 14–10,000 nM were injected for 120 s, followed by 600 s of dissociation. After each cycle, ligand and Ni were removed with 350 mM EDTA (60-s injection, 30 μL/min), and the surface was freshly prepared.

Sensorgrams were double-referenced against the reference flow cell and blank buffer injections. Data were analyzed in Biacore Insight Evaluation Software (Cytiva, version 5.0.18.22102) using a 1:1 Langmuir model to determine association (*k*_a_) and dissociation (*k*_d_) rate constants and equilibrium dissociation constants (*K*_D_). Low ligand capture levels minimized avidity effects; fitted *R*_max_ values were consistent with 1:1 stoichiometry. *k*_a_, *k*_d_, *R*_max_, and the mass transfer coefficient were fitted globally across each concentration series, with the bulk refractive index term fixed at zero. Because His-tagged ligand gradually dissociated from Ni-NTA during each cycle, a linear baseline drift term was fitted locally to each sensorgram. The Ang1/Ang2 comparison (Supplementary Tables 1 and 2) and A14 mutant panel (Tables 1 and 2) were acquired independently, each with parallel wild-type Ang2 measurements. Because capture surfaces were renewed each cycle, rate constants and affinities were compared within, not between, series; SI used values from one series.

### Brønsted analysis of binding kinetics

The analysis follows the Leffler–Brønsted description of transition states^40^, formalized as protein-engineering Φ-value analysis^41^ and applied to protein–protein association^42^. Rate constants are positive, multiplicative quantities better described by log-normal than normal distributions^43^. The equilibrium constant *K*_A_ = *k*_a_/*k*_d_ was calculated within each replicate to preserve pairing. For each substitution, the transition-state penalty was ΔΔ*G*‡ = −*RT* ln(*k*_a__mutant / *k*_a__wild-type), and the overall penalty was ΔΔ*G*_eq_ = −*RT* ln(*K*_A__mutant / *K*_A__wild-type), with *RT* = 2.479 kJ/mol at the SPR temperature (298.15 K). Φ = ΔΔ*G*‡/ΔΔ*G*_eq_ was interpreted as the fraction of the final penalty present at the transition state^41^. For the Brønsted analysis and for the comparisons against the number of side-chain atoms beyond Cβ, each constant was summarized on the natural-logarithmic scale: the mean and standard error of ln *k* were formed across the three replicates, and the mean was back-transformed by exponentiation to give the geometric mean, which is the quantity entering ΔΔ*G*‡, ΔΔ*G*_eq_ and Spearman’s ρ. Uncertainty on ln *k* was not back-transformed into a symmetric ± term, which has no meaning on the original scale; it was carried on the logarithmic scale, where it enters the analysis through the propagation to Φ by Monte Carlo sampling of SE[ln *k*_a_] and SE[ln *k*_d_] (200,000 draws). The rate constants and affinities in Tables 1 and 2, Supplementary Tables 1 and 2, and Figures 2 and 4 are, by contrast, arithmetic means ± SEM of the three determinations in each series; for the A14 mutant panel (Tables 1 and 2) these are the same determinations summarized geometrically above. Because the two summaries differ for a log-normal quantity, the two fold changes quoted for the comparison between proteins with and without an atom beyond Cβ are ratios of geometric means and are not reproducible from the arithmetic means in the tables: association at angulin-3 falls from 3.11 × 10^5^ to 4.17 × 10^3^ M^−1^ s^−1^ (74.8-fold) and dissociation at angulin-1 rises from 2.86 × 10^−3^ to 3.85 × 10^−2^ s^−1^ (13.4-fold), each group value being the geometric mean of the system geometric means it contains. The mutant vs. wild-type fold changes quoted elsewhere derive from the arithmetic means tabulated in Tables 1 and 2 and Supplementary Tables 1 and 2.

### MD simulations

Simulations started from top-ranked AF3 models of wild-type Ang2 with angulin-1 or angulin-3 ECDs, predicted as above. Five position 14 mutants were built from these models using the most probable rotamer in PyMOL 3.1.0’s mutagenesis wizard. Thus, each receptor’s six proteins shared a backbone and interface geometry, differing only at position 14. Free Ang2 comprised the isolated ligand chain from the angulin-1 model with the same substitutions. Twelve complexes (six proteins × two receptors) and six free proteins were simulated.

Simulations used GROMACS 2024.3^44^, the AMBER99SB-ILDN force field^45^, and TIP3P water^46^. Systems were placed in cubic boxes of 11.44 nm for angulin-1 complexes, 13.27 nm for angulin-3 complexes, or 7.26 nm for free Ang2, solvated, and neutralized with Na^+^ counterions, yielding 149,395, 229,122, and 37,201 atoms, respectively. Energy was minimized by steepest descent (maximum 50,000 steps; tolerance, 1,000 kJ/mol/nm), followed by 100-ps NVT and 100-ps NPT equilibration with protein heavy-atom position restraints. Production runs lasted 100 ns with 2-fs steps, LINCS constraints on hydrogen-containing bonds^47^, the Verlet cut-off scheme (*r*coulomb = *r*vdw = 1.0 nm), particle-mesh Ewald electrostatics (0.16-nm grid)^48,49^, V-rescale temperature coupling at 300 K (τ = 0.1 ps)^50^, and C-rescale pressure coupling at 1 bar (τ = 2 ps)^51^. Protein coordinates were saved every 10 ps, yielding 10,001 frames per trajectory; analyses used the protein coordinates only.

Three independent replicas were run per system. Replicas differed only in initial velocities, starting from the same equilibrated structure with fresh Maxwell distributions (gen_vel = yes, gen_seed = −1, continuation = no). This yielded 36 complex and 18 free-protein trajectories, totaling 5.4 μs. Trajectories were rebuilt across periodic boundaries before analysis; replicas, rather than frames, were the analytical units.

### Trajectory analysis

Analyses used MDAnalysis 2.9.0^52^ in Python 3.10 with NumPy and SciPy, sampling every 100 ps (1,001 frames per trajectory). An Ang2 residue contacted the receptor when any of its heavy atoms was within 4.5 Å of a receptor heavy atom. Contact occupancy was the fraction of frames meeting this criterion; formed-to-broken transitions measured intermittency. Y99 occupancy and contacting atom-pair counts in the Results were calculated this way.

Solvent-accessible surface area was calculated every 100 ps using gmx sasa and its double cubic lattice method^53^. The bound-state surface comprised the whole complex, allowing receptor occlusion of the epitope; the free-state surface comprised the isolated ligand. All residue 14 heavy atoms formed the output group. Bound-state exposure was normalized to the protein’s mean residue 14 area across three free-state replicas, accounting for intrinsic side-chain size. Because exposure was intermittent, with zero medians in all twelve systems, we report opening frequency (percentage of frames with any residue 14 exposure) and time-averaged exposure. Both were calculated per replica, then averaged across three replicas per system. Proteins were compared by the absence (Ang2, A14G) or presence (A14C, A14S, A14T, A14V) of side-chain atoms beyond Cβ.

### Steered MD

Eighteen starting conformations per complex were sampled at 20, 35, 50, 65, 80, and 95 ns from each of three replicas. Chains were rebuilt as one cluster, which places both chains in the same periodic image; the chain center-of-mass separation of each conformation was then checked against the bound-complex range of 3.0–5.0 nm, and all 216 conformations passed (observed range, 3.24–3.80 nm). Each conformation was rotated to align the ligand-to-receptor center-of-mass vector with +*z*, placed in a rectangular box with 1.6-nm padding, solvated with TIP3P water and neutralized at 0.15 M NaCl, minimized, and equilibrated for 200 ps in NVT and 1 ns in NPT with position restraints, ensuring identical preparation.

The receptor was pulled from the ligand along *z* using a center-of-mass umbrella potential^54^, a force constant of 1,000 kJ/mol/nm^2^, and a rate of 0.5 nm/ns for 4.5 ns, starting with fresh Maxwell velocities. Pull-coordinate force was saved every 1 ps; separation was complete by approximately 3.2 ns in every run. This yielded 216 pulls, 18 per complex. Peak separation force was the maximum restoring-force magnitude per pull, averaged across six conformations per replica to yield the three system values in Figures 7F and 7G. At this far-from-equilibrium rate, peak force is not a free energy^55^ and is compared only across systems using this protocol^56^.

### Statistical analyses

Cell-based assays were analyzed using GraphPad Prism 9.4.1 (GraphPad Software, Boston, MA, USA). Each well was a separate permeable support carrying its own monolayer, seeded and treated individually, and the well was the experimental unit of the analysis: n is the number of replicate wells per treatment group within a single experiment (3–5, as stated in each figure legend). Bars are means ± standard errors of the mean (SEM) of those wells, and significance was assessed by one-way ANOVA followed by Dunnett’s multiple-comparison test against vehicle. Because all wells in a given panel come from the same passage and the same experiment, this replication is technical rather than biological, and the *P* values describe variation between wells within one experiment rather than between independent experiments. Each experiment was repeated independently at least three times with consistent results, and the figures show one representative experiment.

Kinetic constants and simulation-derived quantities were analyzed in Python 3.10.20 with NumPy 2.2.6 and SciPy 1.15.2. Because side-chain heavy-atom count beyond Cβ is a protein property, replicates were not treated as independent observations of that count. Rankings used Spearman’s ρ across six system means, with exact permutation *p* values calculated from all 720 label permutations. For the tied predictor (0, 0, 1, 1, 2, 2), the smallest attainable two-sided *p* is 16/720 = 0.0222. All tests were two-sided.

### Use of generative AI

During the preparation of this manuscript, the authors used Claude Opus 5 (Anthropic) for two purposes: to improve the clarity and readability of the manuscript text, and to assist in drafting and debugging the Python scripts used for trajectory analysis. All resulting text was reviewed and edited by the authors. All generated code was reviewed, tested against manually verified reference calculations, and validated by the authors. The authors take full responsibility for the content of this manuscript and for the analyses reported herein.

## Acknowledgments

The authors would like to thank Ms. Yuko Fujieda and Mr. Kazuki Matsuo for excellent technical support, and all members of the laboratory for their helpful comments and discussion.

## Author contributions

Conceptualization: T.K. and Y.N. Methodology: T.K. and Y.N. Software: Y.N. Validation: T.K. and Y.N. Formal analysis: T.K. and Y.N. Investigation: T.K., Y.N., and S.K. Data curation: T.K. and Y.N. Visualization: T.K. and Y.N. Supervision: Y.N. and M.K. Project administration: T.K. and Y.N. Funding acquisition: T.K. and M.K. Writing – Original Draft: T.K. and Y.N. Writing – Review & Editing: S.K. and M.K.

## Data availability

All data generated in this study are available in the Zenodo repository under accession code 10.5281/zenodo.24103964 (https://doi.org/10.5281/zenodo.24103964). The deposition comprises the AlphaFold3 predictions, with all diffusion samples and confidence files for every job; the molecular dynamics run inputs for all 18 simulated systems and the steered molecular dynamics run inputs for the 12 complexes, comprising starting coordinates, topologies, force-field include files, position restraints, index files and run parameter files; the protein coordinates of all 54 production trajectories; and the numerical values underlying the figures and tables. The protein sequences used for structure prediction are available from NCBI RefSeq under accessions WP_003463384.1 (*Clostridium perfringens* iota-toxin receptor-binding component Ib), NP_991403.2 (human angulin-1), NP_001186728.1 (human angulin-2) and NP_955383.1 (human angulin-3). No experimentally determined structure of an angubindin–angulin complex is available, and none was used; all structural models reported here are predictions.

## Code availability

The analysis code is available at https://github.com/Yakuji/ang2-a14-analysis under the MIT license and is archived at Zenodo under accession code 10.5281/zenodo.24551832 (https://doi.org/10.5281/zenodo.24551832). The data that the code reads are available in the Zenodo repository under accession code 10.5281/zenodo.24103964.

