## Supplementary document for "Steric bulk at one position of a tight junction modulator switches the kinetic step governing angulin paralog selectivity"

**Supplementary Table 1. SPR-derived kinetic parameters for Ang1 and Ang2 binding to human angulin-1.**

|  | ***k*_a_ (× 10^6^ M^−1^ s^−1^)** | ***k*_d_ (× 10^−3^ s^−1^)** | ***K*_D_ (nM)** |
| --- | --- | --- | --- |
| **Ang1** | 3.07±0.32 | 2.52±0.14 | 0.84±0.08 |
| **Ang2** | 1.41±0.04 | 1.99±0.06 | 1.42±0.03 |

Data are arithmetic means ± SEM from three independent experiments.

**Supplementary Table 2. SPR-derived kinetic parameters for Ang1 and Ang2 binding to human angulin-3.**

|  | ***k*_a_ (× 10^5^ M^−1^ s^−1^)** | ***k*_d_ (× 10^−2^ s^−1^)** | ***K*_D_ (nM)** |
| --- | --- | --- | --- |
| **Ang1** | 1.29±0.08 | 1.490±0.026 | 116.0±8.2 |
| **Ang2** | 2.76±0.64 | 2.323±0.416 | 87.3±6.9 |

Data are arithmetic means ± SEM from three independent experiments.

**Supplementary Table 3. Primers used for site-directed mutagenesis of position 14 of Ang2.**

| **Mutant** | **Forward primer (5′→3′)** | **Reverse primer (5′→3′)** |
| --- | --- | --- |
| A14C | ATATTATTGTGATATAAAGCTTGACAC | ATATCACAATAATATTTGTTATGTGC |
| A14D | ATATTATGATGATATAAAGCTTGACAC | ATATCATCATAATATTTGTTATGTGC |
| A14E | ATATTATGAAGATATAAAGCTTGACAC | ATATCTTCATAATATTTGTTATGTGC |
| A14F | ATATTATTTTGATATAAAGCTTGACAC | ATATCAAAATAATATTTGTTATGTGC |
| A14G | ATATTATGGTGATATAAAGCTTGACAC | ATATCACCATAATATTTGTTATGTGC |
| A14H | ATATTATCATGATATAAAGCTTGACAC | ATATCATGATAATATTTGTTATGTGC |
| A14I | ATATTATATTGATATAAAGCTTGACAC | ATATCAATATAATATTTGTTATGTGC |
| A14K | ATATTATAAAGATATAAAGCTTGACAC | ATATCTTTATAATATTTGTTATGTGC |
| A14L | ATATTATCTGGATATAAAGCTTGACAC | ATATCCAGATAATATTTGTTATGTGC |
| A14M | ATATTATATGGATATAAAGCTTGACAC | ATATCCATATAATATTTGTTATGTGC |
| A14N | ATATTATAATGATATAAAGCTTGACAC | ATATCATTATAATATTTGTTATGTGC |
| A14P | ATATTATCCGGATATAAAGCTTGACAC | ATATCCGGATAATATTTGTTATGTGC |
| A14Q | ATATTATCAGGATATAAAGCTTGACAC | ATATCCTGATAATATTTGTTATGTGC |
| A14R | ATATTATCGTGATATAAAGCTTGACAC | ATATCACGATAATATTTGTTATGTGC |
| A14S | ATATTATAGCGATATAAAGCTTGACAC | ATATCGCTATAATATTTGTTATGTGC |
| A14T | ATATTATACCGATATAAAGCTTGACAC | ATATCGGTATAATATTTGTTATGTGC |
| A14V | ATATTATGTTGATATAAAGCTTGACAC | ATATCAACATAATATTTGTTATGTGC |
| A14W | ATATTATTGGGATATAAAGCTTGACAC | ATATCCCAATAATATTTGTTATGTGC |
| A14Y | ATATTATTATGATATAAAGCTTGACAC | ATATCATAATAATATTTGTTATGTGC |

The substituted codon is underlined in each primer; the reverse primer carries the reverse complement of the same codon. Position 14 of Ang2 corresponds to Ala558 of the mature Ib component (Ang2 position = Ib number − 544).

All plasmids were generated by PCR from pET-16b-Ang2 as template.

**Supplementary Figure legends**

**Supplementary Figure 1. AlphaFold3 confidence across the A14 substitution series is reproducible over independent seeds.**

Ang2 was modeled in complex with the human angulin-1 or angulin-3 ECD for wild-type Ang2 and all 19 nonalanine A14 substitutions, repeating every complex with five independent seeds; each seed returned five diffusion samples. (A) Interface confidence (ipTM). (B) Whole-complex confidence (pTM). Black, angulin-1; red, angulin-3. Columns denote the residue at position 14 in one-letter code; A is wild-type Ang2. Bars are means ± SEM across the five seeds. The receptor asymmetry described in Figure 3B is reproduced under seed resampling: substitutions lower angulin-3 confidence far more than angulin-1 confidence, most steeply at A14D, while angulin-1 ipTM remains high for every substitution.

**Supplementary Figure 2. SPR sensorgrams for wild-type Ang2 and the five A14 mutants.**

His-tagged wild-type Ang2 or the indicated A14 mutant was captured on a Sensor Chip NTA, with human angulin ECD–rabbit Fc fusion proteins as analytes. (A) Angulin-1. (B) Angulin-3. Each panel shows wild-type Ang2 and A14C, A14G, A14S, A14T, and A14V. Colored traces are double-referenced sensorgrams at the analyte concentrations indicated, prepared as a threefold dilution series. One of the three independent determinations summarized in Tables 1 and 2 is shown for each protein.

**Supplementary Figure 3. Baseline TER did not differ across treatment groups in the Caco-2 experiments.**

TER of mature Caco-2 monolayers measured immediately before basal treatment, for the experiments shown in Figure 5. (A–F) The six treatment series of Figure 5: proteins at 50 µg/mL (A), and the concentration responses of A14C (B), A14G (C), A14S (D), A14T (E), and A14V (F). Bars are means ± SEM of replicate wells within a single experiment, each well a separate insert carrying its own monolayer; *n* per group as in Figure 5. Ordinary one-way ANOVA with Dunnett’s multiple-comparison test against vehicle detected no difference in any series.

**Supplementary Figure 4. Baseline TER did not differ across treatment groups in the EpH4 experiments.**

TER of EpH4 monolayers measured immediately before basal treatment, for the experiments shown in Figure 6. (A) Parental EpH4 cells expressing endogenous angulin-1. (B) Angulin-1-KD EpH4 cells. (C) Angulin-2-expressing EpH4 cells. (D) Angulin-3-expressing EpH4 cells. Bars are means ± SEM of replicate wells within a single experiment, each well a separate insert carrying its own monolayer; *n* = 5 per group. Ordinary one-way ANOVA with Dunnett’s multiple-comparison test against vehicle detected no difference in any series.

**Supplementary Figure 5. The rate constants that do not govern selectivity show no ordering by side-chain atom count, and position 14 exposure follows the same receptor contrast.**

(A, B) Rate constants not responsible for paralog selectivity, plotted against side-chain heavy-atom count beyond Cβ (*γ*): association at angulin-1 (A) and dissociation at angulin-3 (B). Three independent SPR determinations per protein; horizontal lines are arithmetic means of the three determinations; *ρ* denotes Spearman’s correlation across six proteins. Neither ordering is monotonic, in contrast to the selectivity-determining constants in Figure 7A and 7B. (C) Time-averaged solvent exposure of position 14 in the complex, expressed as a percentage of its exposure in free Ang2, at angulin-1 (left) and angulin-3 (right). Three independent 100-ns replicas per system; symbols are replica values, horizontal lines are arithmetic means across replicas, and the dotted line separates *γ* = 0 from *γ* = 1 or 2.
