## Supplementary Figure for "Steric bulk at one position of a tight junction modulator switches the kinetic step governing angulin paralog selectivity"

### Slide 1
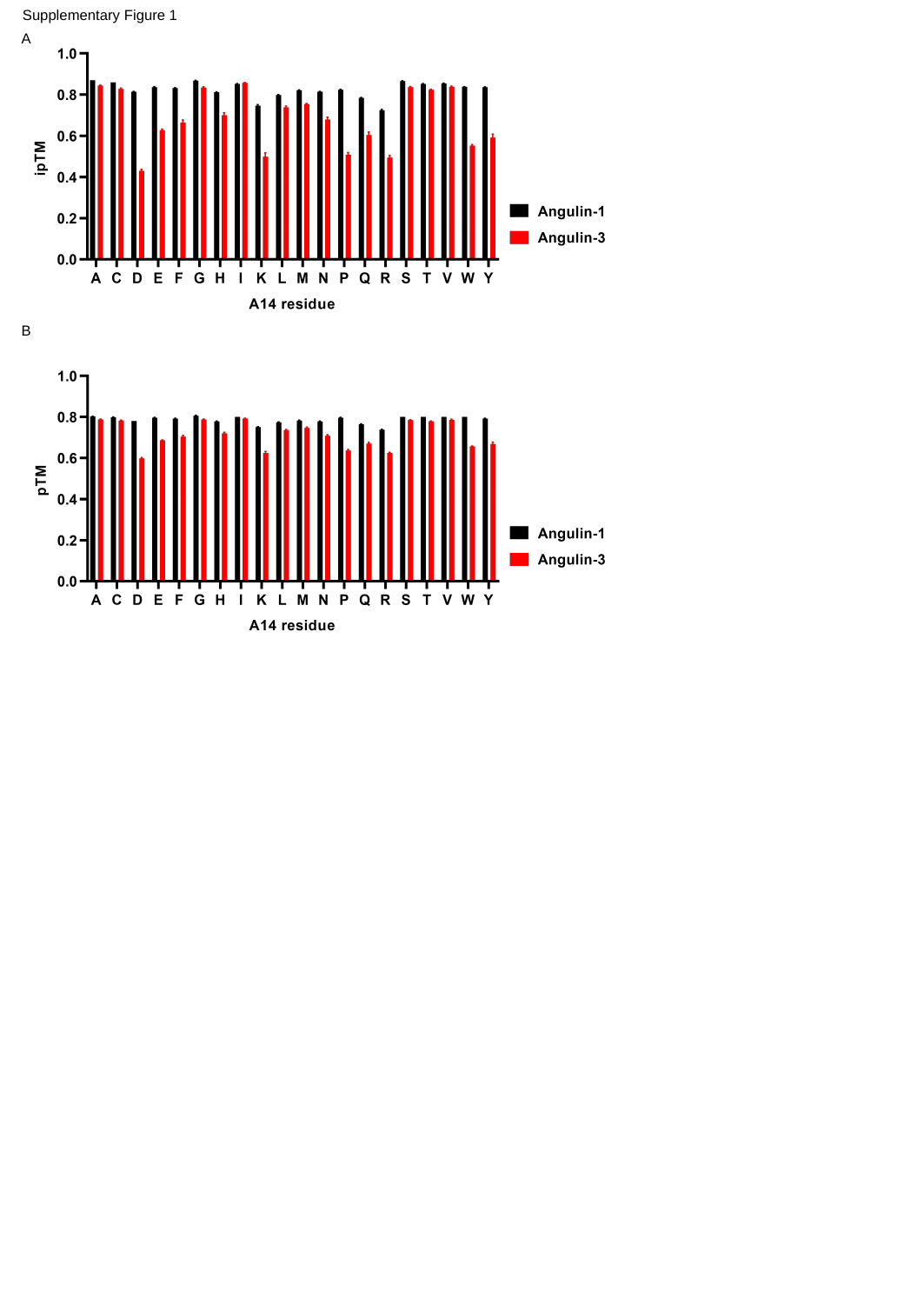

Supplementary Figure 1
A
B

### Slide 2
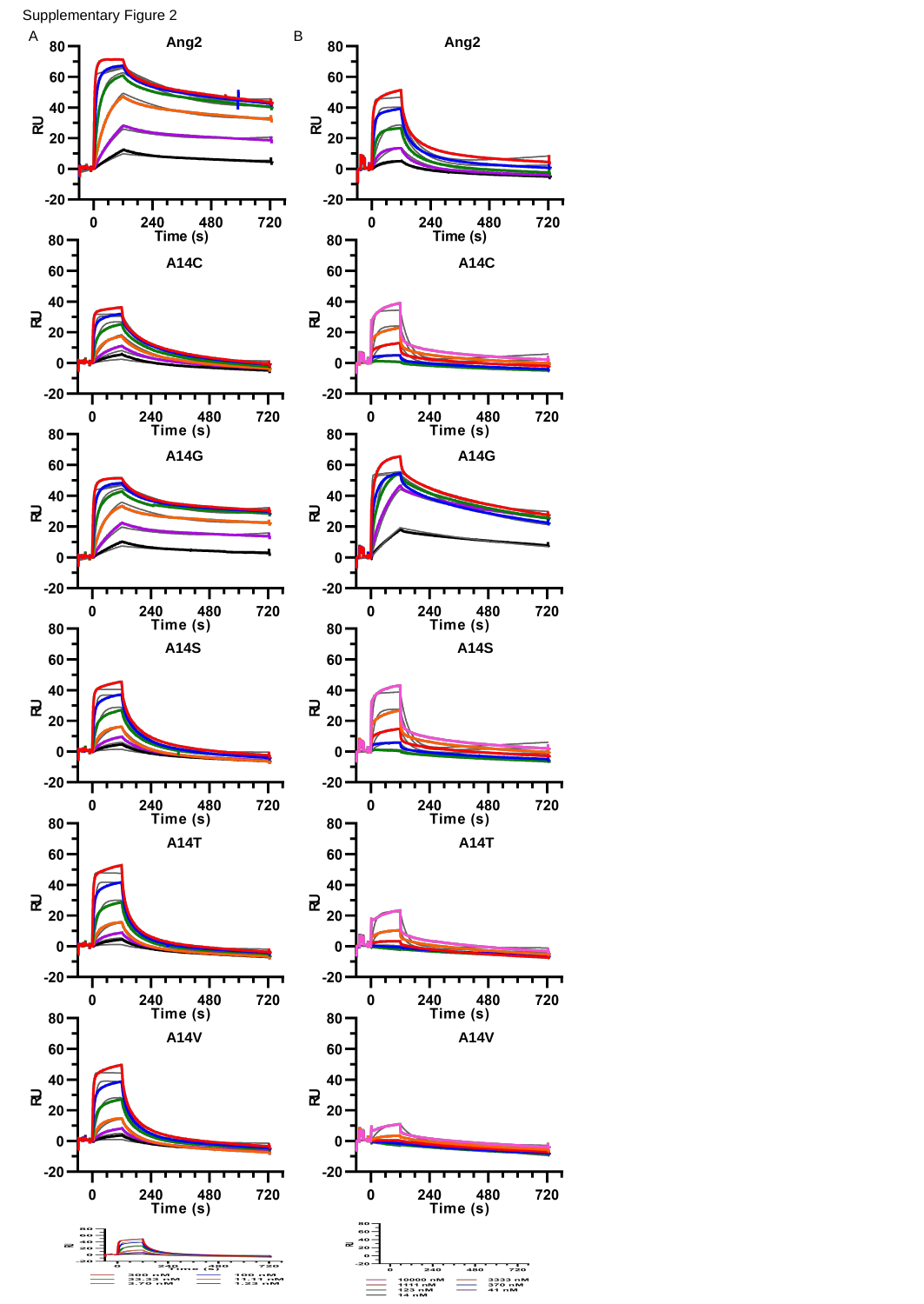

Supplementary Figure 2
A
B
Ang2
Ang2
A14C
A14C
A14G
A14G
A14S
A14S
A14T
A14T
A14V
A14V

### Slide 3
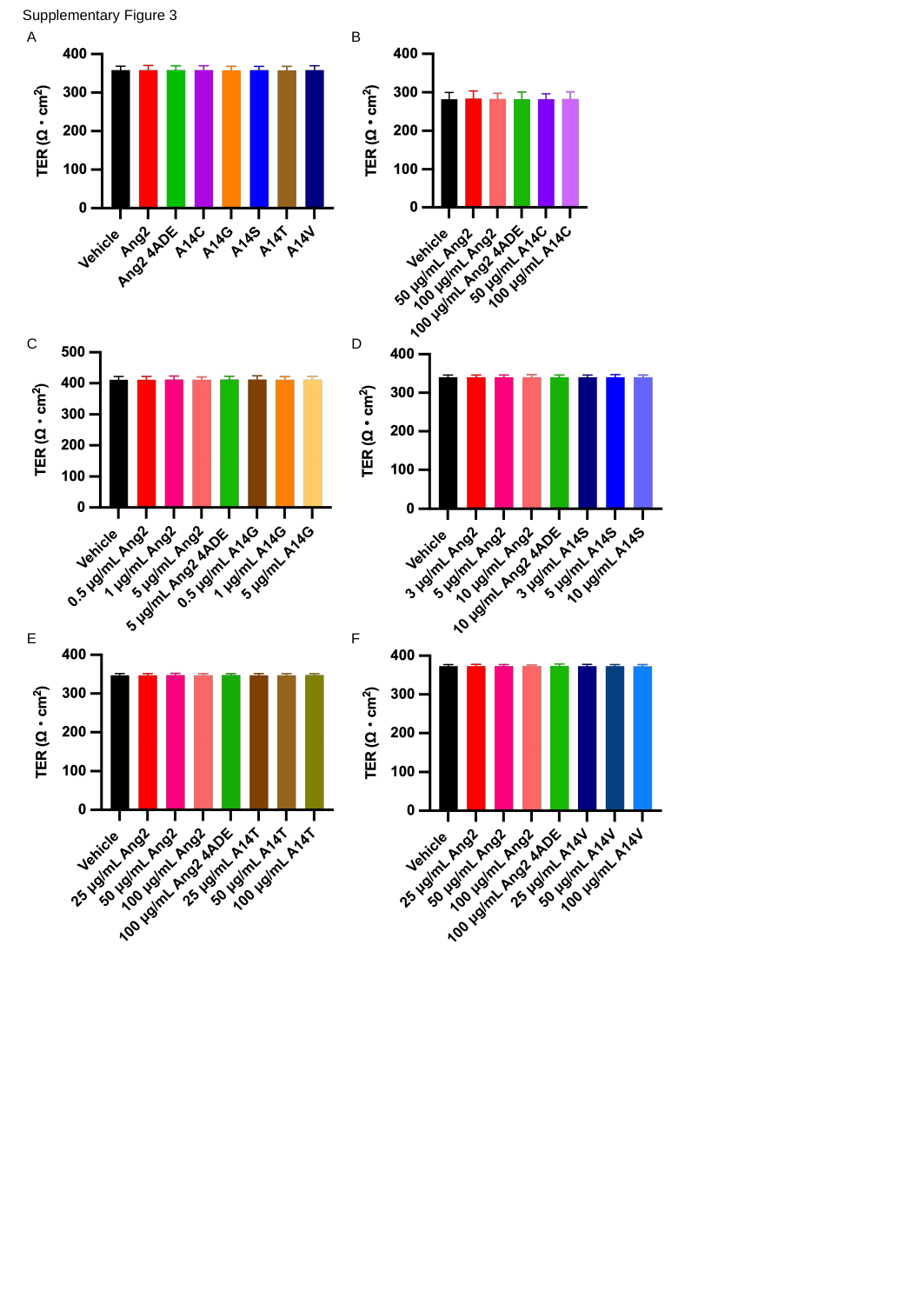

Supplementary Figure 3
A
B
C
D
E
F

### Slide 4
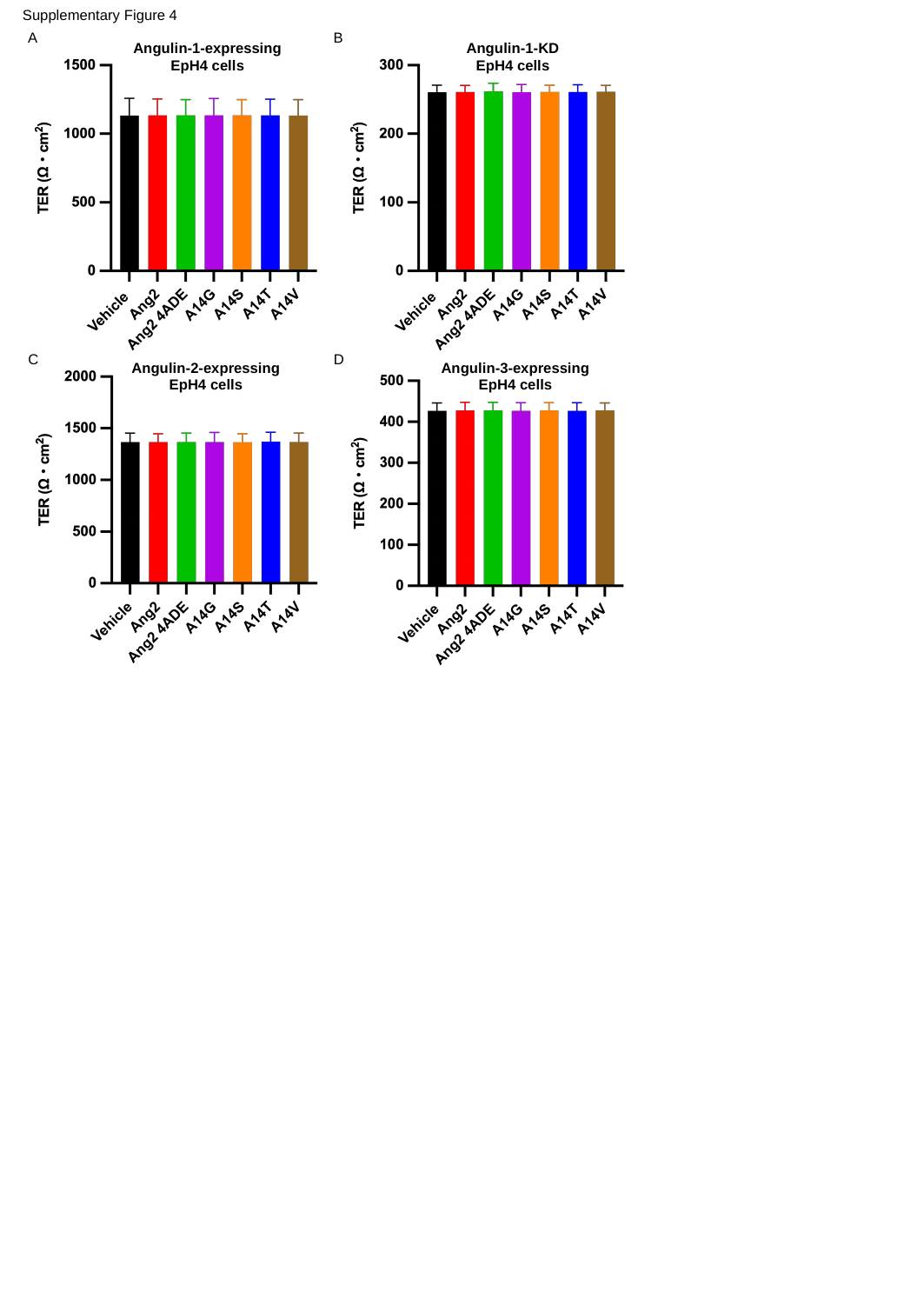

Supplementary Figure 4
A
B
Angulin-1-expressing EpH4 cells
Angulin-1-KD EpH4 cells
C
D
Angulin-2-expressing EpH4 cells
Angulin-3-expressing EpH4 cells

### Slide 5
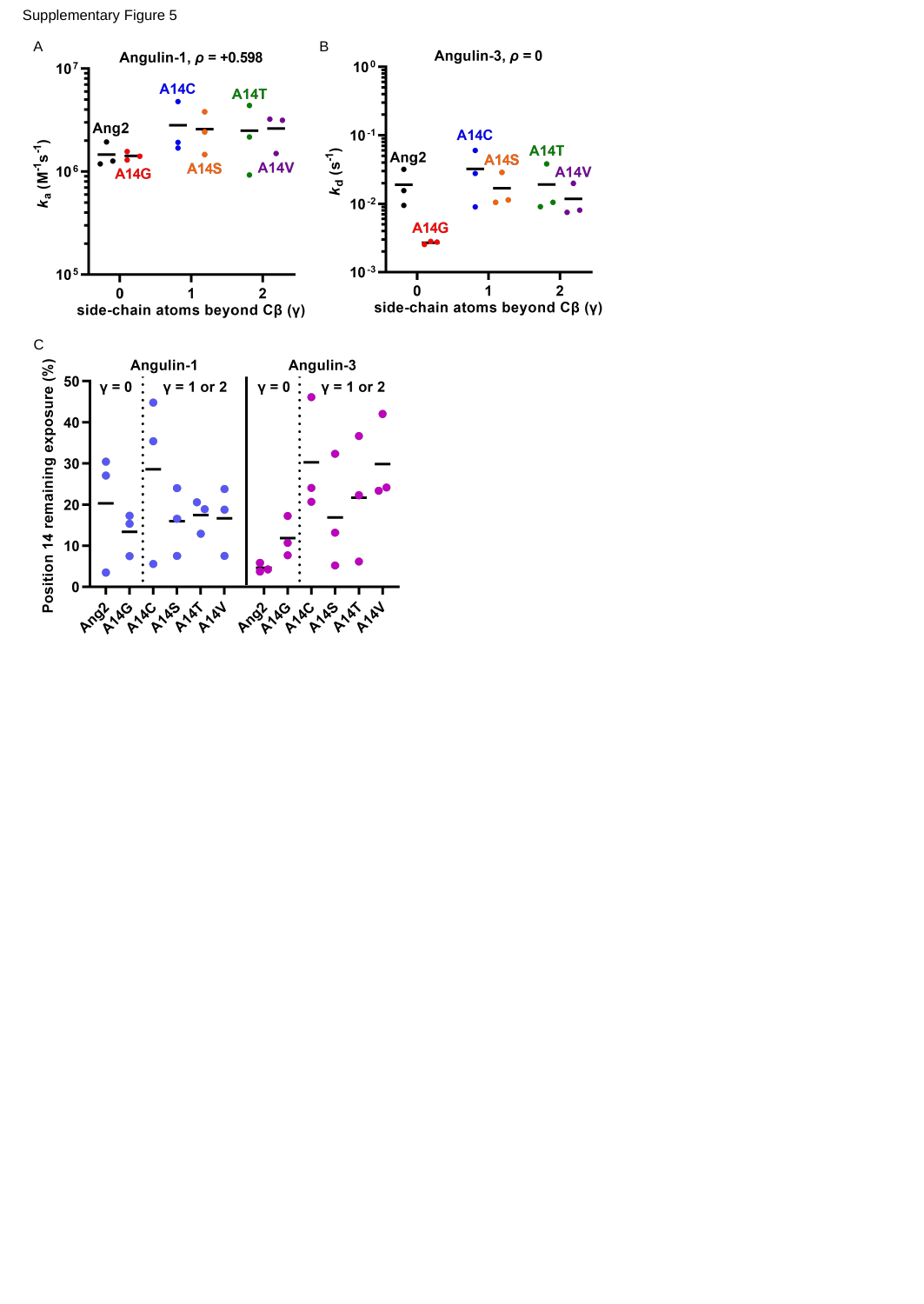

Supplementary Figure 5
A
B
C
